# *Fusobacterium nucleatum* host cell binding and invasion induces IL-8 and CXCL1 secretion that drives colorectal cancer cell migration

**DOI:** 10.1101/2020.01.15.907931

**Authors:** Michael A. Casasanta, Christopher C. Yoo, Barath Udayasuryan, Blake E. Sanders, Ariana Umaña, Yao Zhang, Huaiyao Peng, Alison J. Duncan, Yueying Wang, Liwu Li, Scott S. Verbridge, Daniel J. Slade

**Author notes:** These authors contributed equally (Michael A. Casasanta, Christopher C. Yoo, Barath Udayasuryan). To whom correspondence should be addressed: Dr. Daniel J. Slade, Department of Biochemistry, Virginia Polytechnic Institute and State University, Blacksburg, VA 24061.

## Abstract

*Fusobacterium nucleatum* is implicated in the acceleration of colorectal cancer (CRC), yet the mechanisms by which this bacterium modulates the tumor microenvironment remain understudied. Here we show that binding and cellular invasion of CRC cells selectively induces the secretion of the pro-inflammatory and metastatic cytokines IL-8 and CXCL1, which we then show induces robust migration of HCT116 cancer cells. Next, we demonstrate that cytokine signaling by cancer cells is largely driven by invasion coordinated by the surface adhesin Fap2. By contrast, we show that *F. nucleatum* induced secretion of CCL3, CXCL2, and TNFα cytokines from neutrophils and macrophages is Fap2 independent. Finally, we show that inhibiting *F. nucleatum* host-cell binding and entry using galactose sugars, neutralizing membrane antibodies, and deletion of the *fap2* gene, lead to attenuated cytokine secretion and cellular migration. As elevated IL-8 and CXCL1 levels in cancer have been associated with increased metastatic potential and cell seeding, poor prognosis, and enhanced recruitment of tumor-associated macrophages and fibroblasts within tumor microenvironments, these data show that *F. nucleatum* directly and indirectly modulates immune and cancer cell signaling and migration. In conclusion, as viable *F. nucleatum* were previously shown to migrate within metastatic CRC cells, we propose that inhibition of host cell binding and invasion, potentially through vaccination or novel galactoside compounds, could be an effective strategy for reducing *F. nucleatum*-induced signaling that drives metastasis and cancer cell seeding.

## INTRODUCTION

The role of bacteria and viruses in the onset and progression of diverse cancers is well established (*1–3*). Recent studies on the oral, anaerobic, gram-negative bacterium *Fusobacterium nucleatum* in the acceleration of CRC pathogenesis have revealed as many questions as they have answered for the mechanisms and proteins this bacterium uses to potentiate disease (*4–9*). An overarching question in the field is: How does *F. nucleatum* enter and reside in tumors after likely arriving via the bloodstream from its native oral cavity? In addition, a second theme of pathogenesis that remains understudied is: Are these bacteria capable of leaving the primary tumor on or within immune or cancerous cells to seed and accelerate metastatic cancer sites (*10*)? Answering these questions will be key in understanding both the host and bacterial mechanisms at play in microbe-accelerated cancers. Two recent studies reported that *F. nucleatum* directly induces cancer cell metastasis through NF-κB increased expression of Keratin 7 (KRT7) (*11*), as well as increased expression of caspase activation and recruitment domain 3 (CARD3), and downregulation of E-cadherin (*12*). Herein we add to the mechanisms used by *F. nucleatum* to induce cellular migration. We show that direct binding and invasion of host cancer and immune cells by *F. nucleatum* induces the secretion of the proinflammatory and prometastatic cytokines IL-8 and CXCL1, and that conditioned media from *F. nucleatum* infected HCT116 CRC cells causes non-*Fusobacterium* exposed cells to migrate towards this cytokine rich media.

Chemokines/cytokines play a crucial role in tumor initiation, progression, and metastasis (*13*). Initially discovered as chemotactic mediators of leukocytes, they are now known to be secreted by several cell types and can be expressed constitutively or induced by inflammatory stimuli, including bacterial infections, and function in a variety of roles including cell survival, proliferation, angiogenesis, and cell migration. In cancer, chemokines mainly function in regulating angiogenesis, activating tumor-specific immune responses, and directly stimulating the tumor through autocrine or paracrine mechanisms (*13*).

The cytokines IL-8 (CXCL8) and CXCL1 (GROα) are well known to play a pivotal role in CRC progression. Multiple studies have indicated their important role in influencing CRC invasiveness. Elevated CXCL1 is correlated with cancer progression and metastasis and ultimately poor prognosis in patients with CRC; thus indicating its possible role as a biomarker for CRC (*14*). CXCL1 is an autocrine growth factor that binds to the receptor CXCR2 with high affinity. Many colorectal adenocarcinoma cell lines (LS174T, KM12L4, KM12C, SW480, HT29, and Caco2) constitutively express CXCL1, and it is well known that high levels of this cytokine increase invasive potential (*15*). Additionally, anti-CXCL1 or anti-CXCR2 antibodies have been shown to inhibit colon cancer cell proliferation (*16*). It is relevant for these studies that non-metastatic Caco2 and low-metastatic HT29 cell lines expressed lower levels of CXCL1 than the highly metastatic cancer cell line LS147T. In another study, Ogata et al. showed that CXCL1 increases the number of invasive DLD-1 and LoVo cells, and that these effects are quenched in the presence of anti-CXCL1 antibody (*17*). Furthermore, there is emerging evidence to indicate that CXCL1 participates in premetastatic niche formation in liver tissue which in turn recruits CXCR2-positive myeloid-derived suppressor cells (MDSC) to support liver metastases of CRC (*18*).

IL-8 is a ubiquitously prevalent cytokine in CRC where it has been characterized as the most potent neutrophil chemoattractant and activator in both *in vivo* and *in vitro* studies. Rubie et al. has shown that both IL-8 mRNA and protein expression were significantly upregulated in pathological colorectal tissues as well as enhanced in colorectal liver metastasis (*19*). Chen et al. showed that IL-8 and its receptor CXCR2 were highly upregulated in colon cancer (*20*). Additionally, Lee et al. showed that elevated levels of IL-8 in the serum and tumor microenvironment had enhanced the growth of human and mouse colon cancer cells while promoting the extravasation of cancer cells to lung and liver (*21*). IL-8 can bind to both CXCR1 and CXCR2, but it exerts different effects upon binding to either receptor (*22*). Binding to CXCR1 induces neutrophil migration, whereas binding to CXCR2 modulates angiogenic activity (*13*). The angiogenic effect of IL-8 promotes tumor growth by providing access to oxygen and nutrients, as well as an opportunity to metastasize.

Our initial goal was to investigate the role of outer membrane adhesins in *F. nucleatum* direct binding and invasion of cancer cells to determine if this was critical for altered cell signaling. Multiple adhesins have been characterized in *F. nucleatum* binding and signaling, with important roles for the small multimeric adhesin FadA in E-cadherin interactions and Wnt signaling that drives cellular proliferation (*6, 23*). Additional experiments uncovered the critical role of the large, outer membrane adhesin Fap2, a member of the Type 5a autotransporter protein family (*23–25*). This protein docks with host cells through Gal/GalNAc sugar residues overexpressed on CRC cells, as well as protein-protein interactions with TIGIT on natural killer cells (*26*). However, aside from these two adhesins, most outer membrane proteins of *F. nucleatum* have not been characterized in cancer cell interactions. We have expanded upon these analyses by developing a new, modified version of a galactose kinase markerless gene deletion system capable of creating strains with unlimited gene deletions. We implemented this system to functionally characterize the role of the Type 5a autotransporter adhesin Fap2, multiple uncharacterized Type 5c trimeric autotransporter adhesins (CbpF, FvcB, FvcC, FvcD) (*27, 28*), and the small, non-autotransporter multimeric adhesin FadA. Our studies reveal that binding and invasion of HCT116 CRC cells by *F. nucleatum*, which can be inhibited by small molecules, antibodies, or adhesin gene deletions, is critical to induce pro-inflammatory signaling cascades as well as cellular migration.

Through the results presented here we propose that *F. nucleatum* modulates the tumor microenvironment through interactions with both CRC and immune cells, thereby leading to the secretion of cytokines that have been well characterized for their pro-oncogenic, metastatic, and immune cell recruitment functions. Our results show how a bacterium may accelerate but not initiate cancer, and how these cytokines could allow long-term, low bacterial load infections within tumors to propagate autocrine and paracrine signaling, leading to *F. nucleatum*-loaded cancer cells that initiate cellular seeding at distant sites including the liver.

## RESULTS

### *F. nucleatum* 23726 outer membrane adhesins are critical for the binding and invasion of HCT116 CRC cells

Previous studies have established that *F. nucleatum* is highly invasive and can undergo a non-obligate intracellular life stage within epithelial, endothelial, keratinocytes, and potentially immune cells (*29–32*). We confirm the invasive potential of *F. nucleatum* subsp. *nucleatum* ATCC 23726 into HCT116 CRC cells using fluorescence microscopy (**Fig 1A-C**), imaging flow cytometry (**Fig. 1D**), and classic flow cytometry (**Fig 1E**) to validate our forthcoming experiments evaluating specific proteins in this invasion process.

**Fig. 1.**
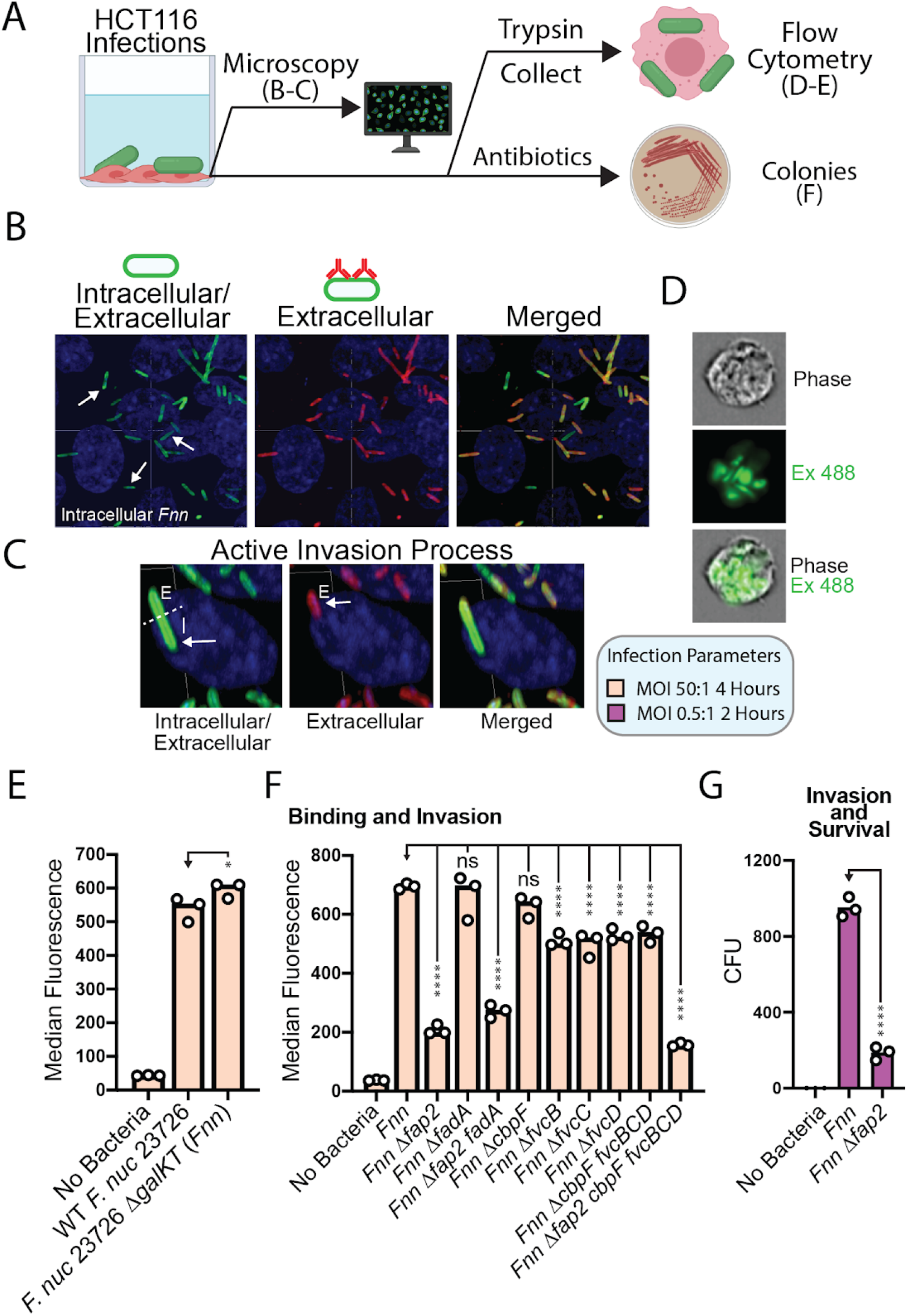
*F. nucleatum* binds to and invades HCT116 cells. (**A**) Overview of experiments used to analyze binding and invasion of *Fnn* and adhesin mutants in HCT116 CRC cells. (**B**) Intra- and extracellular *Fnn* detected with a fluorescent lipid dye (Green, intra- and extracellular) and a pan-*Fusobacterium* membrane antisera (Red, extracellular), host cell nuclei are stained with DAPI (blue). (**C**) Zoomed in view and focus on a single *Fnn* bacterium that is half intracellular, half extracellular. (**D**) Imaging flow cytometry view of intracellular *Fnn*. (**E**) Comparison of HCT116 binding and invasion of wild type (WT) *F. nucleatum* 23726 with *F. nucleatum* 23726 Δ*galKT* (*Fnn*). (**F**) Binding and invasion analysis of *Fnn* and adhesin gene deletion strains using flow cytometry. (**G**) Invasion and survival of *Fnn* and *Fnn* Δ*fap2* in HCT116 cells analyzed using an antibiotic protection assay.

To gain a deeper understanding of how *F. nucleatum* is contributing to the acceleration of cancer, we set out to understand the importance of verified cancer cell interactions, which are driven by surface exposed outer membrane adhesins. Our goal was to confirm the roles of FadA and Fap2 in HCT116 binding and signaling, as well as perform the first characterization of multiple Type 5c trimeric autotransporter adhesins which have a well-established role in the virulence potential of other gram-negative bacteria such as *Yersinia (33, 34*). We recently bioinformatically identified five Type 5c adhesins in the strain *F. nucleatum* 23726, with four of these (CbpF, FvcB, FvcC, FvcD) containing all of the classic domains that make up a complete adhesin. The fifth protein (FvcE, not characterized) lacks a ‘head’ domain that is predicted to coordinate adhesion (*23*). To study these proteins, we made the first complete, single gene knockouts of these six adhesin genes (*fadA, fap2, cbpF fvcB, fvcC, fvcD*) in *F. nucleatum* 23726, as well as multiple gene deletions per strain (**Table S1–3**) using a new version of a galactose kinase (GalK) genetic system previously used in *F. nucleatum* (*35*), as well as many classical studies in *Clostridium (36*). The details of developing this efficient genetic system and gene deletions used in this study can be found in **Figures S1–6** and the *Materials and Methods.*

We first established that the deletion of *galKT,* which remains in all of the subsequent mutant strains, did not affect growth (**Fig. S3H**) and binding to HCT116 cells (**Fig. 1E**) when compared to wild type (WT) *F. nucleatum* 23726. We have used the nomenclature *Fnn* to describe the *F. nucleatum* 23726 Δ*galKT* strain throughout the paper and figures (**Fig. 1E**).

We next show that *Fnn* Δ*fap2* is significantly attenuated for HCT116 cellular interactions (**Fig 1F**), but that *Fnn* Δ*fadA* shows no decrease in HCT116 interactions. While FadA has been shown to drive interactions with cancer cells, we note that these studies were done in the strain *F. nucleatum* 12230 (*31, 37, 38*). Upon bioinformatic analysis, we found that *F. nucleatum* 23726 contains four *fadA* ortholog genes (*fadA2, fadA3a, fadA3b, fadA3c*), and *F. nucleatum* 12230 encodes for only one orthologue (*23*). This could mean that a single deletion of the *fadA* gene in *F. nucleatum* 23726 remains adhesive, and that multiple *fadA* family proteins are contributing to binding and invasion. In addition, a *Fnn* Δ*fap2 fadA* double mutant did not further reduce binding when compared to *Fnn* Δ*fap2* (**Fig. 1F**). Concurrently, we analyzed single mutants of the trimeric autotransporter adhesins *cbpF, f vcB, fvcC, fvcD*, as well as a strain lacking all four genes (*Fnn* Δ*cbpf fvcBCD*). We show that FvcB, FvcC, and FvcD contribute to invasion, albeit less potently than Fap2. Importantly, a *Fnn* Δ*fap2 cbpf fvcBCD* quintuple mutant showed no significant decrease in HCT116 invasion when compared to the single *fap2* gene deletion strain. Last, we show that the proportion of intracellular *Fnn* and *Fnn* Δfap2 determined by antibiotic protection assays (**Fig. 1G**) correlates with the flow cytometry data, indicating that the trypsinization used to prepare infected HCT116 cells likely removes the majority of surface bound bacteria. Taken together, these data indicate that host cell binding and invasion is largely driven by Fap2.

### F. nucleatum host-cell binding and invasion is critical for inducing the secretion of pro-inflammatory and pro-metastatic cytokines from cancer cells

After confirming the importance of outer membrane adhesins in invading HCT116 cells, we analyzed the ability of this bacterium to induce the secretion of cytokines from both human cancer cells and mouse neutrophils and macrophages. Using cytokine arrays (**Fig. 2A-C**), we report that *F. nucleatum* 23726 interactions with HCT116 cells induces the secretion of high concentrations (~ 1000 pg/mL after four hours) of the CXC family cytokines IL-8 and CXCL1 into the culture medium (**Fig. 2D**). To determine if direct binding and invasion is necessary for induced cytokine secretion, as well as to rule out whether an unidentified secreted protein or molecule induces this phenomenon, we compared *Fnn* to *Fnn* Δ*fap2*. Our results show that the ability of *Fnn* Δ*fap2* to induce cytokine secretion from HCT116 cells is significantly attenuated. This correlates well with reduced invasive potential and confirms direct interaction is needed to alter host cells (**Fig. 2D**).

**Fig. 2.**
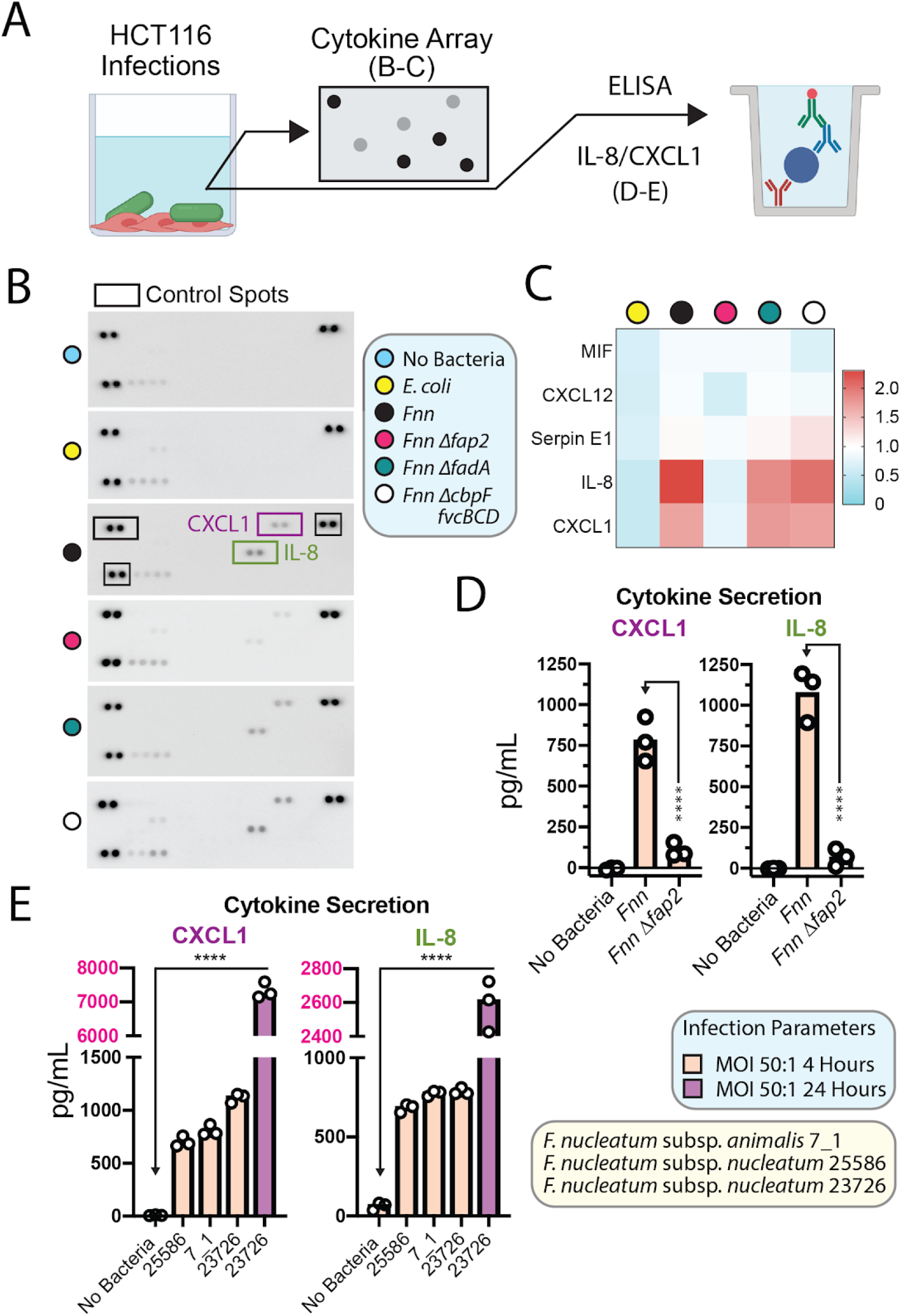
*F. nucleatum* induces cytokine secretion from HCT116 cells. (**A**) Overview of experiments used to analyze *Fnn* induced cytokine secretion from HCT116 CRC cells. (**B**) Broad cytokine array dot blots analyzing *Fnn* and *Fnn* adhesin deletion strains. (**C**) Heatmap of fold increased dot blot intensity reveals an increase in IL-8 and CXCL1 secretion for invasive strains. (**D**) IL-8 and CXCL1 ELISA to quantitate cytokine secretion from HCT116 cells induced by *Fnn* and *Fnn* Δ*fap2.* (**E**) IL-8 and CXCL1 ELISA to quantitate and compare cytokine secretion from HCT116 cells induced by *F. nucleatum* subsp. *nucleatum* 25586, *F. nucleatum* subsp. *nucleatum* 23726, *F. nucleatum* subsp. *animalis* 7_1 (*Fna*).

To test if this phenomenon was specific to strain *F. nucleatum* 23726, we compared IL-8 and CXCL1 secretion levels to *F. nucleatum* 25586 and 7_1, which cover subspecies *nucleatum* and *animalis* (*Fna*) respectively. We show that these strains also induce comparable, significant levels of cytokine secretion. Lastly, we compared our 4 hour secretion levels to that of the 24 hour level for *F. nucleatum* 23726 and report an increase in CXCL1 and IL-8 to ~ 7000 pg/mL and ~2500 pg/mL respectively.

### *F. nucleatum* induced cytokine secretion in neutrophils and macrophages is not Fap2 dependent

We characterized cytokine secretion from mouse immune cells using a mouse-specific cytokine array. We first concurrently tested neutrophil cytokine secretion (**Fig. 3B**), as well as validated the direct binding and potential invasion of neutrophils via fluorescence microscopy (**Fig. 3C**). *F. nucleatum* induces the secretion of CCL3, CXCL2, and TNFα, and we validate high amounts of CCL3 and CXCL2 from neutrophils via ELISA (**Fig. 3D**). In mouse macrophages, *F. nucleatum* also induces CCL3, CXCL2, and TNFα secretion (**Fig. 3E**), as well as CCL5 and additional cytokines when compared to either no bacteria or *E. coli* control infections. CCL3 has a defined role in lymphocyte recruitment in early metastatic CRC (*39*), and CXCL2 has been shown to promote angiogenesis in CRC tumor microenvironments (*40*). Interestingly, microscopy shows a smeared DNA pattern from infected neutrophils, potentially indicating the formation of neutrophil extracellular traps (NETs) and a pathway for cytokine secretion (*41*). *F. nucleatum* and other oral bacteria were previously shown to degrade NETs, which is a potential escape mechanism from neutrophil-mediated death (*42*). In stark contrast to the Fap2-driven cytokine secretion seen in HCT116 cancer cells, *Fnn* Δ*fap2* did not show decreased levels of CCL3 or CXCL2 secretion as analyzed by cytokine arrays and ELISA. We conclude that this shows the importance of Fap2 in selective cancer cell binding, which is evidenced by a lack of Gal/GalNAc sugar on immune cells. In addition, we acknowledge that a role of immune cells is to engulf and clear invading organisms like bacteria, and therefore these cells have acquired the ability to recognize a broad range of bacterial ligands as endocytosis targets.

**Fig. 3.**
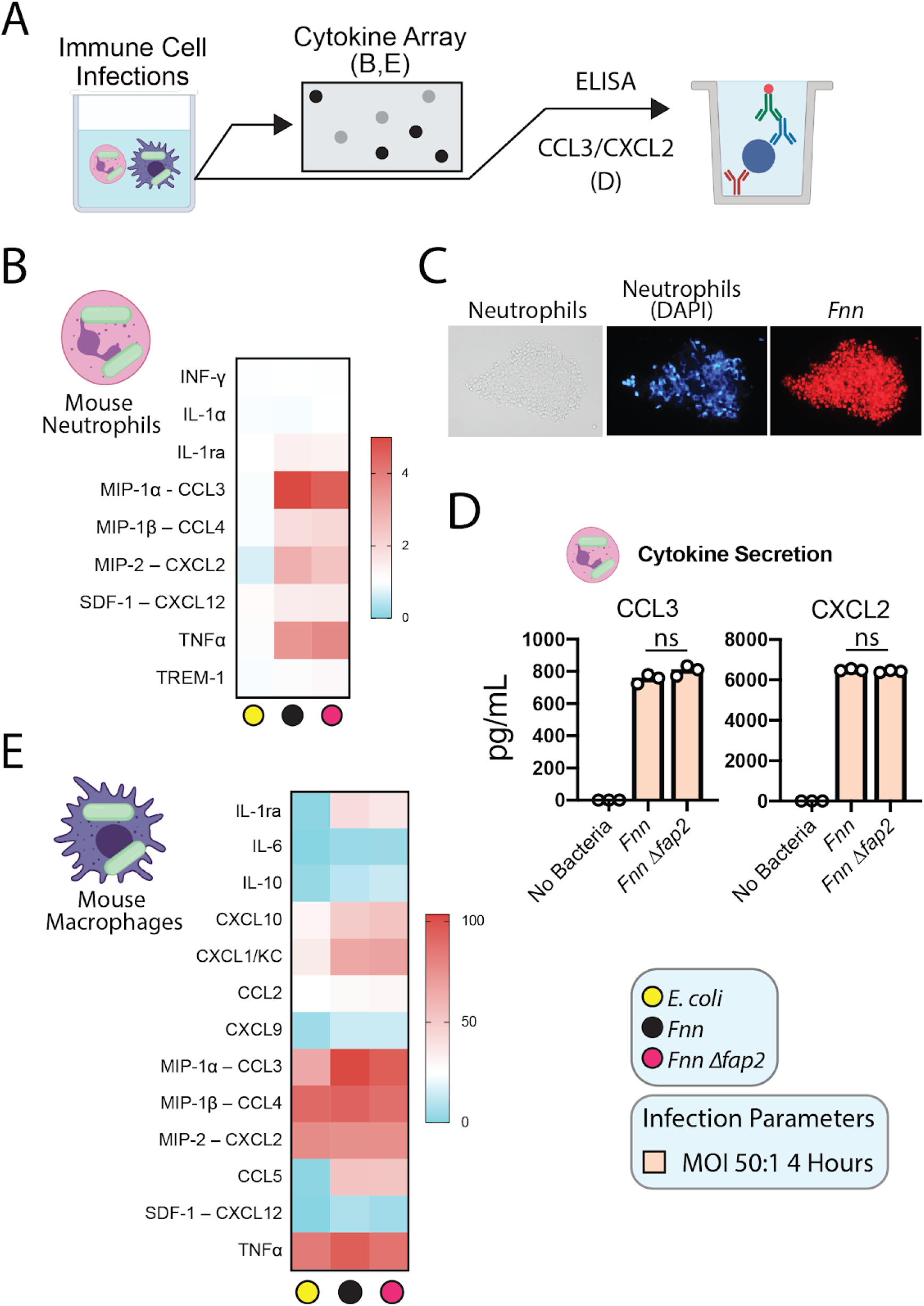
Cytokine secretion analysis from mouse neutrophils and macrophages. (**A**) Overview of experiments used to analyze *Fnn* induced cytokine secretion from immune cells. (**B**) Mouse cytokine array shows *Fnn* induced secretion of CCL3, CXCL2, and TNFα from neutrophils. (**D**) Fluorescence microscopy of *Fnn* interacting with mouse neutrophils. (**D**) ELISA confirming *Fnn* induction of CCL3 and CXCL2, and revealing that deletion of *fap2* does not decrease cytokine secretion as it does in HCT116 cells. (**E**) Mouse cytokine array shows *Fnn* induced secretion of several cytokines from macrophages when compared to *E. coli* and no bacteria controls.

### Inhibition of bacterial invasion using sugars and arginine blocks cytokine secretion

Since Fap2 is a Gal/GalNAc sugar binding lectin, and this sugar is overrepresented on the surface of CRC cells, we next tested a panel of galactose and non-galactose containing sugars for their ability to inhibit HCT116 binding and IL-8 and CXCL1 secretion. We show that galactose, GalNAc, and lactose (galactose disaccharide) potently inhibit HCT116 binding and invasion, while control sugars including glucose and maltose do not significantly inhibit invasion (**Fig. 4A-B**). In addition, we tested L-arginine as an inhibitor because the *F. nucleatum* surface adhesin RadD, which is predominantly thought to coordinate interspecies bacterial interactions in the oral biofilm, is an L-arginine inhibitable adhesin (*43–45*). We show that L-arginine inhibits HCT116 cellular invasion, but not to the extent of galactose sugars. All sugar molecules tested and L-arginine lead to reduced IL-8 and CXCL1 secretion (**Fig. 4C**), with the most potent being GalNAc as seen in the binding and invasion assay. We believe the addition of 10 mM glucose and maltose results in non-specific inhibition of cytokine secretion, or could be binding to an unidentified *F. nucleatum* lectin.

**Fig 4.**
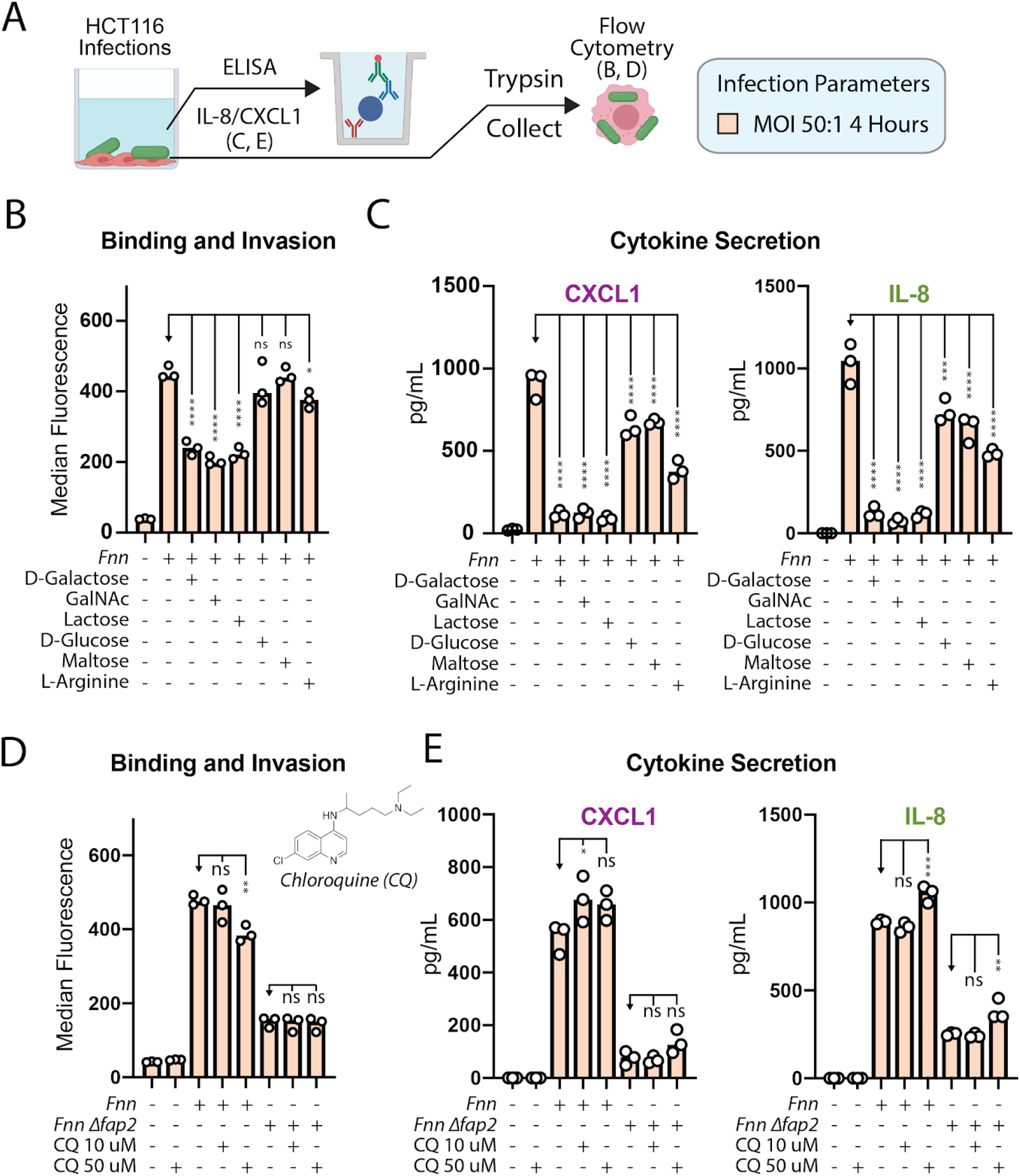
Inhibition of *Fnn* HCT116 binding and invasion using small molecules. (**A**) Overview of experiments used to analyze small molecule inhibitors of *Fnn* binding and invasion of HCT116 CRC cells. (**B**) *Fnn* invasion is significantly inhibited by galactose containing sugars and L-arginine (10 mM). (**C**) Secretion of IL-8 and CXCL1 from HCT116 cells is inhibited by all sugars tested and L-arginine (10 mM). (**D**) *Fnn* invasion is not inhibited by 10 μM chloroquine (autophagy and endosome maturation inhibitor), and (**E**) to the contrary could increase CXCL1 and IL-8 secretion.

Finally, we tested the autophagy and endosome maturation inhibitor chloroquine for its ability to inhibit invasion and signaling (**Fig. 4D**). This experiment was performed because the two previous studies reporting *F. nucleatum* induced metastasis showed chloroquine is able to inhibit cellular migration, and we set out to test if this is due to reduced cytokine levels (*11, 12*). We show that chloroquine does not inhibit bacterial invasion at 10 μM, and slightly decreases invasion into HCT116 cells at 50 μM. In addition, we saw the reverse effect of chloroquine on cytokine signaling, with a slight increase in secreted IL-8 and CXCL1. We elaborate on this finding in the *Discussion*.

### IL-8 and CXCL1 induce HCT116 cellular migration

Based on previous reports of IL-8 induced migration of CRC cells (*46*), endothelial cells, and leukocytes (*47*), as well as a role for CXCL1 induced metastasis in multiple cancers including colorectal (*13, 18, 48*), we tested their roles in HCT116 migration using transwell assays (**Fig. 5A-C**). We show that IL-8 and CXCL1 separately induce robust HCT116 migration at 100 ng/mL (**Fig. 5D**), but when added together at the same concentration do not additively increase migration.

**Fig. 5.**
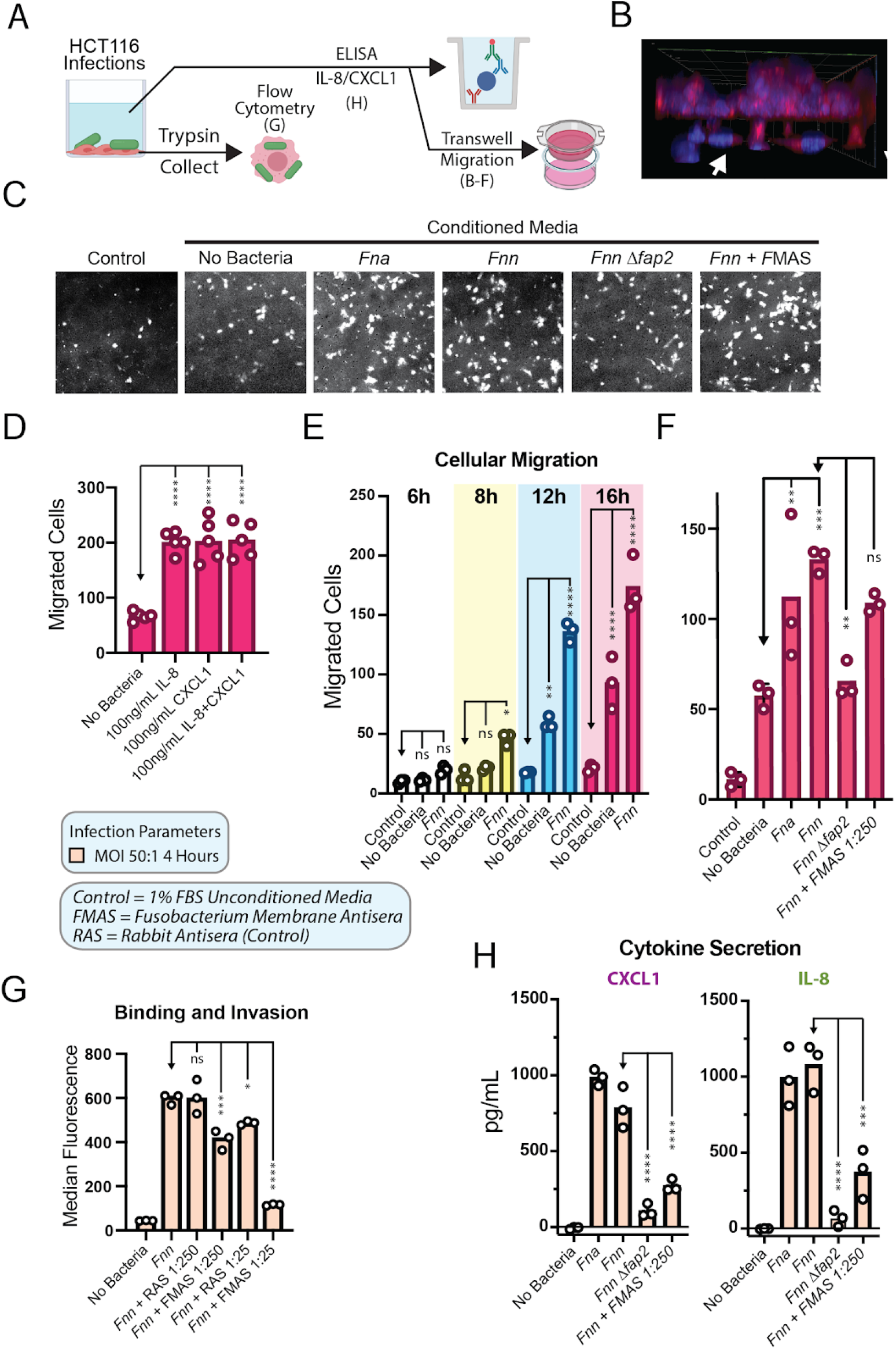
HCT116 migration is driven by *Fnn*-induced secretion of CXCL1 and IL-8. (**A**) Experimental setup of HCT116 transwell migration experiments. (**B**) 3D confocal image of HCT cells (blue and red) migrated across the transwell barrier (white arrows). (**C**) Representative images of migrated HCT cells after exposure to the indicated conditioned media. (**D**) The addition of IL-8 and CXCL1 to culture media within the lower transwell chamber leads to significant HCT116 cancer cell migration. (**E**) HCT116 cellular migration induced by *Fnn* conditioned media with high CXCL1 and IL-8 concentrations. (**F**) Inhibition of HCT cellular migration through deletion of *fap2 (Fnn Δfap2*) and *Fusobacterium* membrane antisera (FMAS). (**G**) Inhibition of HCT116 binding and invasion by *Fnn* using FMAS and control rabbit antisera (RAS). (**H**) Inhibition of CXCL1 and IL-8 secretion from HCT116 cells using FMAS.

### *F. nucleatum* conditioned supernatants from HCT116 infections cause cancer cell migration

After showing that purified IL-8 and CXCL1 induce cancer cell migration, we tested if *F. nucleatum* conditioned culture media from HCT116 infections could cause non-infected cells to migrate. We show through transwell assays that *F. nucleatum* conditioned HCT116 culture media significantly induces non-*Fusobacterium* exposed cancer cells to migrate starting at 8 hours, with increasing significance through our 16 hour test (**Fig. 5E**). We subsequently demonstrate that infection with *Fnn* Δ*fap2,* which leads to lower levels of IL-8 and CXCL1 secretion, results in significantly reduced cell migration when compared to *Fnn* conditioned media (**Fig. 5F**).

We developed a pan-Fusobacterium membrane antisera (DJSVT_MAS1) to potentially block HCT116 cell migration. However, our results show only a marginal reduction in cellular migration. This could be due to insufficient antibody concentrations added to the assay, or since the anti-sera was developed with 11 strains of *Fusobacterium*) its specific inhibitory effects on *F. nucleatum* 23726 have been distributed across non-*Fnn* proteins. However, the pan-*Fusobacterium* membrane antisera was able to significantly decrease cancer cell binding (**Fig. 5G**) and cytokine secretion (**Fig. 5H**).

Finally, we show that bead-conjugated antibody depletion of IL-8 and CXCL1 from *F. nucleatum* conditioned culture media significantly reduces the amount of migrated cells (**Fig. 6**), confirming our results with purified cytokines and conditioned media that IL-8 and CXCL1 specifically drive HCT116 cellular migration. As the depletion of these cytokines does not completely abolish *Fnn* conditioned media induced cell migration, we hypothesize that *F. nucleatum* modulates multiple host proteins and pathways involved in cellular migration and metastasis. We elaborate on this in the Discussion.

**Fig. 6.**
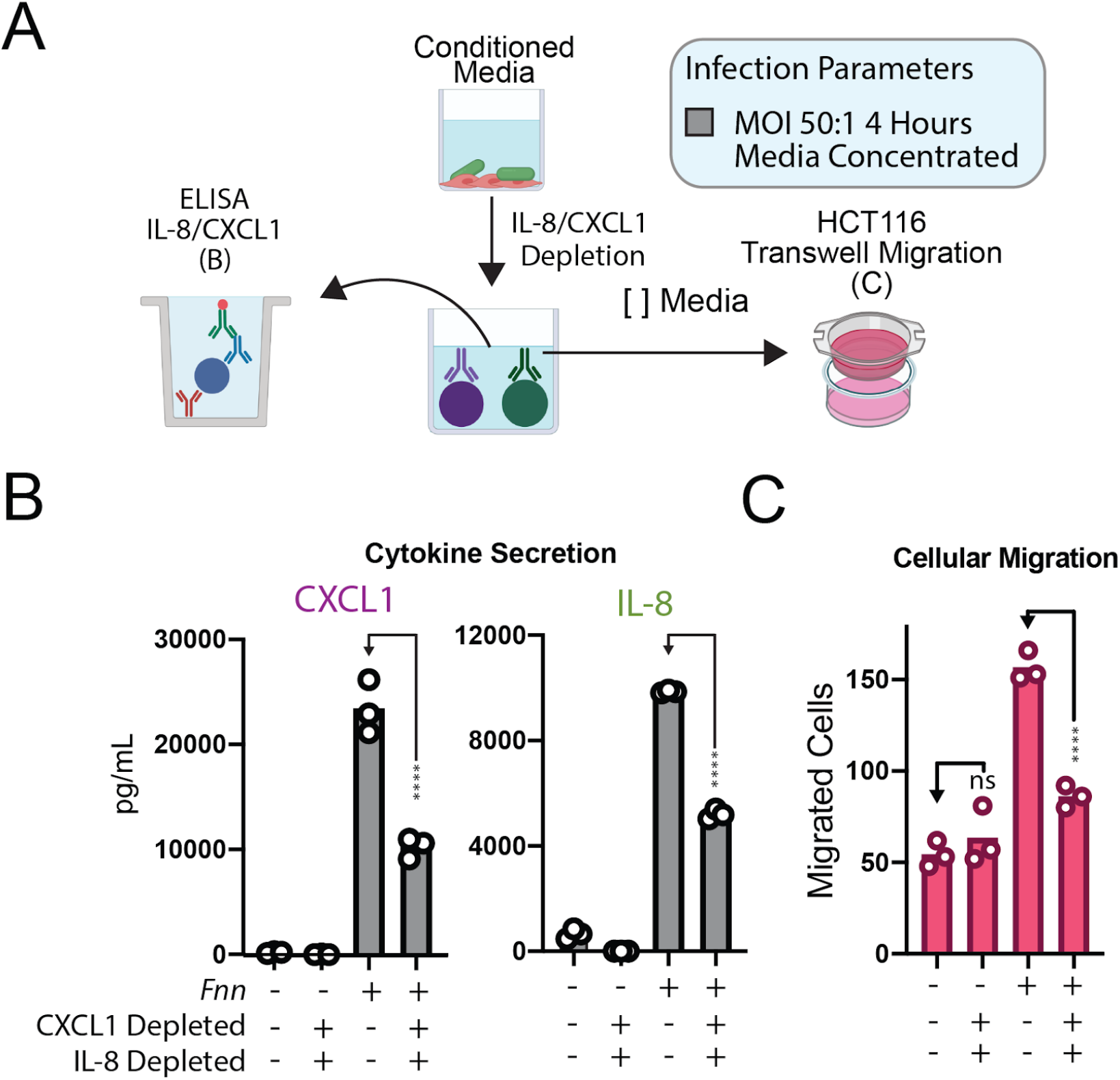
Depletion of CXCL1 and IL-8 from *Fnn* conditioned media block cancer cell migration. **(A)** Experimental setup of HCT116 transwell migration experiments including cytokine depletion from conditioned media. **(B)** Analysis of CXCL1 and IL-8 levels after concentration, with and without antibody depletion of cytokines and before loading into the bottom chamber of the transwell. **(C)** Cellular migration is significantly decreased by depleting CXCL1 and IL-8, nearly to the level of non-*Fusobacterium* conditioned media even through cytokines were not completely depleted.

## DISCUSSION

Studies have illuminated a strong correlation between the presence of *Fusobacterium nucleatum* within colorectal tumors and deleterious patient prognosis, which is proposed to be causally related to increased tumor microenvironment inflammation, enhanced oncogenic signaling (*49–52*). To address the role of *F. nucleatum* in tumor microenvironment signaling, we set out to characterize potential signaling pathways that drive cellular migration and immune cell-driven cancer acceleration. Our data show that direct *F. nucleatum-cellular* interactions with cancer and immune cells, largely coordinated by the outer membrane adhesin Fap2, drive host cells to define a locally pro-metastatic, inflammatory microenvironment. The ability of this bacterium to cause selective induction of the proinflammatory and prometastatic cytokines IL-8 and CXCL1 from CRC cells complements prior studies in which these gross phenotypes of oncogenic acceleration from human patients and mouse models are consistently reported (*10, 53, 54*). In addition, our findings add to a growing understanding that *F. nucleatum* directly induces metastasis through cytokine release, increased NF-κB expression and subsequent expression of KRT7 (*11*), increased CARD3, and downregulation of E-cadherin (*12*). These studies have now shown active roles for *F. nucleatum* in mouse models of metastasis, which complement the seminal study by Bullman et al. first reporting viable *F. nucleatum* within human CRC cell metastases in the liver (**Fig. 7**) (*10*).

**Fig. 7.**
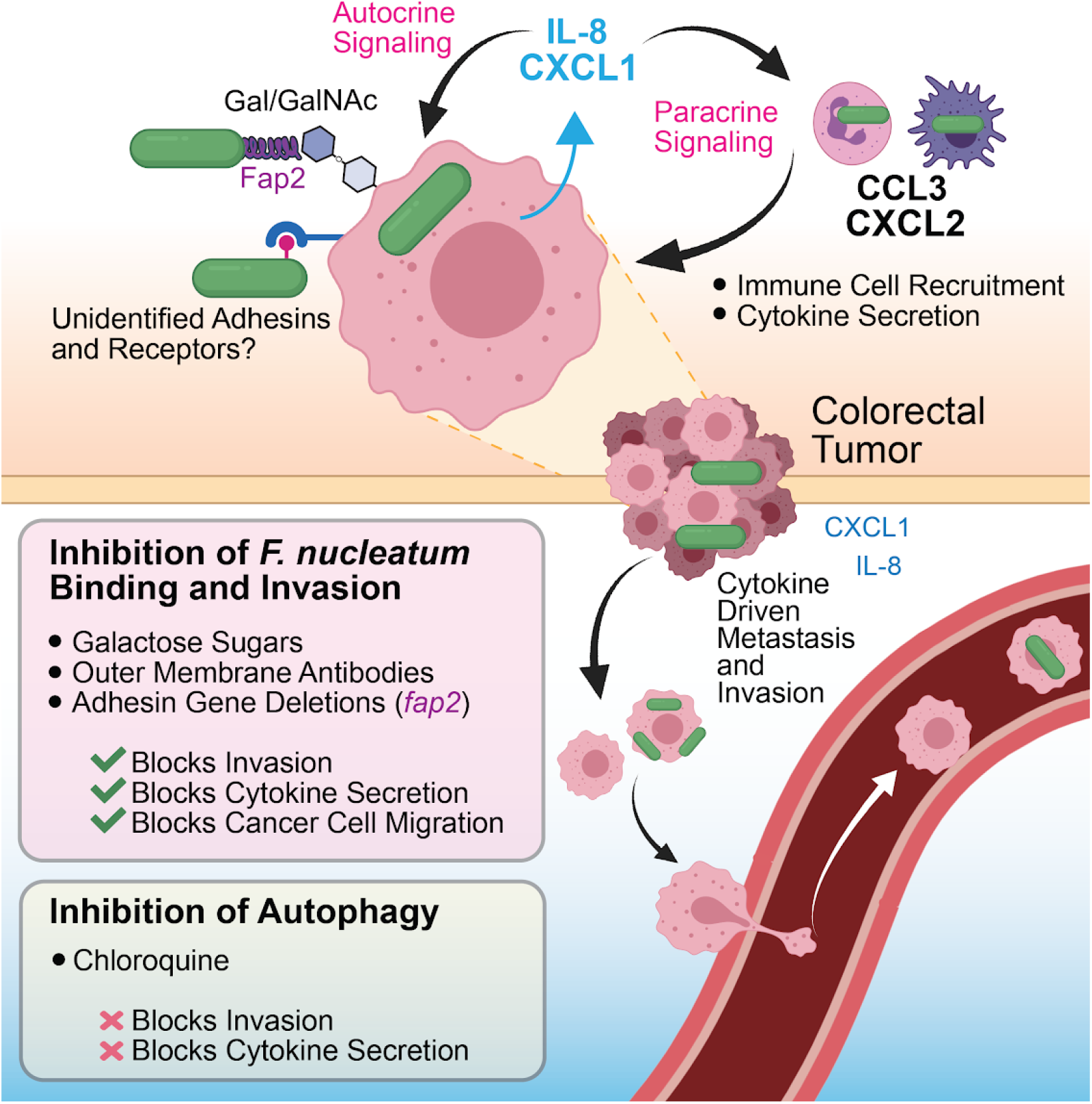
A model of *F. nucleatum* induced metastasis through cytokine signaling. We show that *F. nucleatum* induces IL-8 and CXCL1 secretion from CRC cells, and both cytokines have been characterized as key players in cancer metastasis and subsequent downstream cell seeding. As well as autocrine signaling back to cancer cells as a metastatic signal, HCT116 derived cytokines can participate in paracrine signaling to recruit neighboring immune cells, which further secrete their own cytokine signatures that alter the tumor microenvironment through metastatic, inflammatory, and immune cell programming.

CXCL1 and IL-8 both play roles in immune cell recruitment, particularly neutrophil recruitment and programming. Through paracrine signaling, *F. nucleatum* interactions with CRC cells could be releasing factors that create not only a metastatic environment, but one that provides pro-tumor inflammation. There is the potential for recruited neutrophils to be activated to tumor-associated neutrophils (TAN) that could further accelerate tumor progression. In addition, if *F. nucleatum* or the tumor microenvironment then induces neutrophil extracellular trap (NET) formation, these structures have been shown to increase metastasis and sequester circulating tumor cells to promote reseeding (*55–57*). Finally, it was shown that high intra-tumor loads of *F. nucleatum* resulted lower CD3+ T-cell density in CRC, providing another layer to the complexity of *F. nucleatum* immune cell regulation (*58*). To address these potential scenarios, a priority should be placed on investigating the potential for this bacterium to induce the formation of additional pro-oncogenic cell types including cancer-associated fibroblasts (CAF) (*59*), tumor-associated macrophages (TAM) (*60*), and tumor-associated neutrophils (TAN) (*55*).

Bacteria such as *Streptococcus gallolyticus* are known to induce the secretion of IL-8 in colon tumor cells (*61*). It has also been shown that increased expression of IL-8 leads to a significant resistance to the cytotoxic effects of oxaliplatin, a platinum based chemotherapeutic drug (*46*). As *F. nucleatum* was previously shown to induce chemoresistance through the induction of autophagy and subsequent inhibition of apoptosis (*51*), our report of the induction of IL-8 and CXCL1 by this bacterium adds another dimension to this story.

Our data definitively shows the importance of *F. nucleatum* binding and invasion of human cancer cells in producing pro-metastatic cytokines. We and others have proposed that the development of small molecule compounds or proteins that are able to block *F. nucleatum* docking to host cells could be effective strategies to reduce cancer and immune cell signaling that accelerates metastasis from the colon. A starting point could be the selective inhibition of Fap2, which we have now shown drives binding and cytokine secretion. Fap2 was previously characterized as a lectin and shown to bind to Gal/GalNac sugars that are in abundance on the surface of most cancer cells, especially colorectal (*62*). The potential for targeting of bacterial lectins for disease treatment has been validated in urinary tract infections by using mannoside sugar derivative compounds that block the uropathogenic *E. coli* lectin FimH from binding to host receptors, thereby resulting in depletion of this pathogenic bacteria from the gut (*63*).

Two recent studies report that chloroquine blocks metastasis by lowering *F. nucleatum-induced* NF-κB driven expression of Keratin 7; a protein linked to increased cancer cell motility and invasive potential (*64, 65*), and overexpression of CARD3; a pro-metastatic kinase previously characterized in breast cancer (*66*). We were therefore interested in whether the IL-8 and CXCL1-regulated cell migration we have observed might be mitigated by chloroquine administration. Our results show that chloroquine treatment actually increased the secretion of IL-8 and CXCL1, suggesting that multiple pathways may be at play in regulating *F. nucleatum*-driven metastasis. This outcome is also the opposite of the effect chloroquine has on reducing the amount of IL-8 secreted upon *Campylobacter jejuni* infections (*67*). Taken together these observations could mean the IL-8 and CXCL1 secretion we observed is not caused by a Toll-Like Receptor (TLR) maturation dependent mechanism, for which chloroquine has been shown to regulate (*67, 68*), but is still NF-κB dependent. In addition, inhibiting autophagy could lead to increased intracellular survival of *F. nucleatum*. This should be characterized further before these inhibitors are considered for use as more than chemical tools to understand host biological pathways.

As a final point, since the consensus belief is that *F. nucleatum* leaves the oral cavity and traverses the human body through the blood and potentially lymph, it could be advantageous to use a vaccine-based strategy whereby antibodies block and clear this bacterium prior to leaving the bloodstream, thereby preventing downstream tumor interactions. Our data using a pan-*F. usobacterium* membrane antisera show effective blocking of *F. nucleatum* entry into cancer cells, but this strategy needs refinement to increase strain specific potency. Hence, interfering with *F. nucleatum* interactions in the human body could be a targeted approach that provides an alternative to using non-specific antibiotics. Finally, we hypothesize that future studies characterizing how *F. nucleatum* intricately influences multiple cell signaling pathways during cancer progression and metastasis will lead to targeted approaches for controlling additional oncomicrobes in cancer.

## MATERIALS AND METHODS

### Culturing *Fusobacterium nucleatum*

*Fusobacterium nucleatum* subsp. *nucleatum* ATCC 23726, *Fusobacterium nucleatum* subsp. *nucleatum* ATCC 25586, and *Fusobacterium nucleatum* subsp. *animalis* 7_1 (*Fna*) were cultured on solid agar plates made with Columbia Broth (Gibco) substituted with hemin (5 μg/mL) and menadione (0.5 μg/mL) (CBHK) under anaerobic conditions (90% N_2_, 5% H_2_, 5% CO_2_) at 37 °C. Liquid cultures started from single colonies were grown in CBHK media under the same conditions. For infections, overnight cultures were back diluted 1:1000 and grown to mid-exponential phase (~0.5 OD_600_) prior to subsequent experiments unless otherwise noted.

### Culturing HCT116 CRC cells

HCT116 (ATCC CCL-247) cells were purchased from ATCC and grown on tissue culture treated plates and flasks in McCoy’s 5A media supplemented with 10% fetal bovine serum (FBS), penicillin, and streptomycin. Cells were grown to no more than passage 15 at 37 °C with 5% CO_2_. For maintenance between infections, cells were passaged by gentle trypsinization and reseeding. For infection experiments, cells were grown to >90% confluency in 6 or 24 well plates. Unless otherwise indicated, all experiments were conducted with HCT116 media supplemented with 10% FBS.

### Molecular cloning of a galactose selectable gene deletion system

We have developed a new iteration of a galactose-selectable gene deletion system in *Fusobacterium nucleatum* ATCC 23726 (*Fnn*) capable of deleting an unlimited number of genes in a single strain. We showcase the power of our genetic system by developing strains containing multiple gene deletions from ~300bp-12kb, as well as chromosomally complementing tagged versions of these virulence factors for detection and purification. The plasmids used for these genetic mutations are all new, developed in house, and represented in **Table S2**. This system is different than a previously reported system in that it relies on the deletion of both the galactose kinase (*galK*) and galactose-1-phosphate uridylyltransferase (*galT*) in the Leloir pathway (**Fig. S1)**; ultimately creating a system that allows for the selection of target gene deletions on solid media containing galactose and not the more expensive 2-Deoxy-D-galactose. The first step in creating the chloramphenicol and thiamphenicol selectable plasmid pDJSVT1 was to PCR amplify the *catP* gene from a pJIR750 backbone (*Clostridium* shuttle vector) followed by using overlap extension PCR (OLE-PCR) to fuse a pMB1 *E. coli* origin of replication from pUC19 (**Fig. S2A**)(See **Table S1** for primers). This linear product, which contains a multiple cloning site with GC rich DNA restriction sites (XhoI, NotI, KpnI, and MluI) that are optimized for cloning in DNA from AT rich bacteria such as *Fusobacterium* (~75% AT), was cut with NotI and ligated to circularize the plasmid. This final high copy pDJSVT1 construct is small (~1800 bp), which likely enhances transformation efficiency.

pDJSVT1 is the base vector to create multiple additional vectors to complete the genetic system. To delete the *galK* and *galT* genes in *F. nucleatum 23726* to make the base strain DJSVT02 (**Fig. S3A**), 1000 bp directly up and downstream of the galKT gene cluster were amplified separately, then fused using OLE-PCR. This 2kb PCR product was then digested with KpnI/MluI and ligated into pDJSVT1 digested with the same enzymes. This final vector is pDJSVT13. This vector was electroporated (1-3 μg DNA, 2.5 kV, 50 uF capacitance, 360 OHMS resistance) into competent *F. nucleatum* 23726 and selection on chloramphenicol (single chromosomal crossover), followed by selection on solid media containing 1% 2-Deoxy-D-galactose to select for either *galKT* gene deletions, as the absence of the *galT* gene makes 2-Deoxy-D-galactose non-toxic to *F. nucleatum* (**Fig. S1**).

The next plasmid is for targeted gene deletion in the *F. nucleatum* 23726 ΔgalKT (*Fnn:* strain DJSVT02) background. This vector, pDJSVT7 (**Fig. S2B**), contains a FLAG::galK gene under the control of a *Fusobacterium necrophorum* promoter. Transformation allows for initial chromosomal integration and selection with thiamphenicol, followed by selection for double crossover gene deletions on solid media containing 3% galactose. For this study, this vector was used to create gene deletions in *fap2, fadA, cbpF, fvcB, frcC, and fvcD* in *F. nucleatum* 23726 Δ*galKT.* As shown in Figure S4, 750 bp directly up and downstream of a target gene for intra-bacteria homologous recombination were amplified by PCR, making complementary fragments fused by OLE-PCR. This product (Fig S4A) is ligated into pDJSVT7 using KpnI/MluI restriction sites. This vector is then electroporated (2.5 kV, 50 uF capacitance, 360 OHMS resistance) into competent *F. nucleatum* 23726 ΔgalKT and selected on chloramphenicol (single chromosomal crossover), followed by selection on solid media containing 3% galactose which produces either complete gene deletions or wild-type bacteria revertants. Gene deletions are verified by PCR and sequencing, and we show this system has been accurate down to the single base level (**Fig. S5**). For creating multiple gene deletions in a single strain (**Table S3**), additional vectors are created and mutants made in sequential fashion, as gene deletion leaves no trace of vectors, including excision of the chloramphenicol and galactose selection genes.

The third vector in the suite, pDJSVT11 (**Fig. S2C**)(not used in this study), is to create single copy chromosomal complementations at a static chromosomal location within the *arsB* gene, which is only necessary during times of high arsenic exposure and therefore doesn’t change the phenotype of *Fnn* under any conditions tested (**Fig. 1E, Fig. S3H**).

### RNA Extraction and RT-PCR

*F. nucleatum* cultures were grown to stationary phase and pelleted by high-speed centrifugation (12,000 g for 2 min at room temperature). TRIzol Extraction Isolation of total RNA was performed following manufacturer’s instructions (Invitrogen). Briefly, cell pellets were resuspended in 1 mL of TRIol reagent (Invitrogen) and 0.2 mL of chloroform was added. Solution was centrifuged for 15 minutes at 12,000 g at 4 °C. The RNA-containing aqueous phase was collected and the RNA precipitated after 500 uL of isopropanol had been added. The RNA pellet was then washed with 75% ethanol and centrifuged at 10,000 g for 5 minutes at 4°C. After drying at room temperature for 10 min, the RNA pellet was resuspended in 30 uL sterile RNAse-free water, and solubilized by incubating in water bath at 55 °C for 10 minutes. Total RNA was quantified using Qubit™ RNA HS Assay Kit (ThermoFisher).

Before reverse transcriptase (RT)-PCR, RNA samples were subjected to DNAse treatment. Briefly, 500 ng of total RNA was incubated with DNAse I (Invitrogen) for 2 hours at 37 °C. Following treatment, DNAse I was inactivated using EDTA and heating mixture for 5 minutes at 65 °C. (RT)-PCR was performed using the Takara PrimeScript™ One Step RT-PCR Kit according to the manufacturer’s instructions. The PCR conditions consisted of reverse transcription for 30 min at 50°C, initial denaturation for 2 min at 94 °C, followed by 30 cycles (30 sec at 94°C, 30 sec at 50-62°C, and 30 sec at 68°C) and elongation at 68 °C for 1 min. The expected bands around 250 bp was confirmed on a 1.5% agarose gel. Specific primers to detect knockout of gene and validate intact genes upstream and downstream of gene of interest (Supplementary Table 1) were used to amplify from RNA extracts.

### Development of a pan-*Fusobacterium* outer membrane antisera

To test if broadly-neutralizing polyclonal antisera against *Fusobacterium* outer membrane proteins could be effective at blocking cellular binding and entry, rabbits were injected (New England Peptide) with purified total membrane proteins from 11 strains of *Fusobacterium* that span 7 species (*F. nucleatum* subsp. *nucleatum* 23726, *F. nucleatum* subsp. *animalis* 7_1, *F. nucleatum* subsp. *polymorphum* 10953, *F. nucleatum* subsp. *vincentii* 49256, *F. periodonticum* 2_1_31, *F. varium* 27725, *F. ulcerans* 49185, *F. mortiferum* 9817, *F. gonidiaformans* 25563, *F. necrophorum* subsp. *necrophorum* 25286, *F. necrophorum* subsp. *funduliforme* 1_1_36S). 20 mL of each strain was grown as previously described to stationary phase before pooling. 220 mL of pooled *Fusobacterium* culture was centrifuged and the pellet resuspended in 40 mL of phosphate-buffered saline pH 7.4 (PBS) with 2 roche EDTA free protease inhibitor tablets, frozen at −80 °C, and passed through an Emulsiflux C3 at >15k PSI to lyse cells. Lysate was centrifuged at 18000g for 25 minutes to pellet debris and unlysed cells. The supernatant was carefully transferred to ultracentrifuge tubes, and centrifuged in a 50-Ti rotor at 38,000 rpm (~144kG average) for 1 hr and 40 min. The supernatant was discarded, the membrane protein pellet was gently washed with PBS, and 0.5 mL of PBS with 1% BOG was added to each of the two tubes and gently resuspended by stirring overnight at 4 °C. The protein concentration was determined by BCA prior to shipment to New England Peptide for inoculation into rabbits. The resulting membrane antisera is herein named DJSVT_MAS1, but for clarity is described in Figure legends as FMAS for *Fusobacterium* membrane antisera.

### Immunofluorescence sample preparation using pan-*Fusobacterium* membrane antisera

Confluent HCT116 cells on slides were infected with FM-143FX labeled (Green fluorescent lipid) *F. nucleatum 23726* in wells containing McCoy’s 5A media with 10% FBS and no antibiotics at MOI of 50:1 for four hours at 37 °C with 5% CO_2_. Post infection, media was removed and thoroughly washed with PBS with gentle orbital shaking. Infected cells were then fixed with PBS/3.2% paraformaldehyde for 20 minutes at room temperature, followed by washing with PBS. Cells were blocked with Sea Block (Abcam) for 2 hours at 37 °C, followed by the addition of 1:100 dilution of DJSVT_MAS1 overnight at 4 °C. Slides were washed in PBS and incubated for 1 hour with Alexa Fluor 594 goat anti-rabbit antibody diluted in Sea Block. After washing with PBS, cells were permeabilized with PBS/1% Triton X-100 for 20 minutes. Post-permeabilization, cells were washed in PBS and labeled with DAPI for 30 minutes. Following three final PBS washes, coverslips were mounted with 80% glycerol/0.1M Tris pH 8.5 and sealed onto glass slides. Fluorescence microscopy was performed on a Zeiss LSM 800 confocal microscope.

### Flow cytometry

Mid-log phase *F. nucleatum* were incubated with 5 ug/mL FM 1-43FX Lipophilic Styryl Dye to stain outer membranes and allow for green fluorescence detection. After washing in PBS, bacteria were resuspended in PBS at 100x concentration (MOI 50:1) before adding to cultures of HCT116 in media containing 10% FBS. Infections lasted four hours prior to removing the culture media (in many cases used for ELISAs as described), cells washed twice with PBS, followed by cell recovery using 0.05% trypsin. After neutralizing trypsin with media containing 10% FBS, cells were pelleted and resuspended in PBS containing 20 mM EDTA. Cells were loaded onto a Guava easyCyte 5 flow cytometer (Luminex) and 10,000 cells were collected using an initial single cell gate to measure the median green fluorescence induced by intracellular *F. nucleatum* labeled with FM 1-43FX. Post data acquisition, FlowJo 10 software was used to further refine gating to single cells as well as determine median fluorescence of all samples. FlowJo analysis was then transferred to GraphPad Prism for statistical analysis and figure generation. In **Figure 1D** we show intracellular *F. nucleatum* 23726 using imaging flow cytometry on an Amnis ImageStream X Mk II. For this experiment, the same protocol for bacterial and HCT116 growth, FM 1-43FX labeling, and infection times were used.

### Invasion and survival antibiotic protection assays

HCT116 monolayers were washed one time with PBS and the cells were incubated in media containing 10% FBS and no antibiotics. HCT116 cells were then infected at an MOI of 0.5:1 with exponential phase *F. nucleatum* for 2 hrs at 37 °C with 5% CO_2_. After infecting cells for 2 hrs, the cell culture media was aspirated-off and cells were washed 2 times with antibiotic-containing cell culture media to remove any unbound bacteria. Cells were then incubated in media containing penicillin and streptomycin (readily kills *F. nucleatum* 23726) for 1 hr to kill any remaining extracellular bacteria. At the end of the incubation period, cells were washed twice with PBS. Epithelial cells were then incubated with warm sterile water to lyse HCT116 cells. Lysates were plated on CBHK plates and incubated at 37 °C under anaerobic conditions for 48 hrs followed by colony counting.

### Inhibition of *F. nucleatum*:HCT116 binding and signaling using chemical compounds and antibodies

*F. nucleatum* invasion and induced host cell signaling inhibition by chemical compounds and neutralizing antibodies was analyzed using the standard infection conditions described above. Just prior to adding bacteria, with no pre-incubation time, 10 mM of D-galactose, GalNAc, lactose, D-glucose, maltose, or L-arginine were added to the culture media. For DJSVT_MAS1 *Fusobacterium* membrane antisera, sera was added from between 1:25 to 1:250 dilutions per infection. Chloroquine was added to HCT116 cells at 10 μM for four hours prior to infection, and remained in the culture media during infections. For all compounds, after a four hour infection at MOI 50:1, binding and invasion was analyzed using flow cytometry.

### Isolation of mouse neutrophils and macrophages

Mouse bone marrow neutrophils were isolated from wild type C57BL/6 mice over a 62.5% percoll gradient with centrifugation at 1100g for 30 min. Purity was >90% as determined by flow cytometry analysis with Ly6G+CD11b+ staining. Neutrophils were used immediately after isolation. For mouse bone marrow derived macrophages, bone marrow cells were cultured for 5 days in RPMI 1640 medium supplemented with 10% FBS, 2mM L-glutamine, 1% penicillin/streptomycin, and 10ng/mL M-CSF. Fresh medium was replaced every other day. Wild type C57BL/6 mice were bred and maintained in the animal facility at Virginia Tech in accordance with the Institutional Animal Care and Use Committee (IACUC)-approved protocol.

### Human/Mouse Inflammatory Cytokine Arrays

Exponential phase *F. nucleatum* 23726 or *F. nucleatum* 23726 Δ*fap2* were used to infect HCT116 cell monolayers (100% confluency 1.2 x 10^6^ cells/well for 6 well plate) in antibiotic free cell culture media for four hours at a multiplicity of infection of 50 bacteria per human cell (MOI: 50:1). For neutrophils and macrophages, 1 x 10^6^ cells in suspension were used. After four hours all media (1.5 mL) was collected and sterile filtered through 0.2 μm filters (Millipore Sigma) to remove and bacterial and human cells from the sample. The cytokine array membranes (R&D: Proteome Profiler Human Cytokine Array (ARY005B), Proteome Profiler Mouse Cytokine Array Kit, Panel A (ARY006)) were then blocked for one hour in TBS 3% bovine serum albumin (BSA) at room temperature on a rocking platform. While the membrane was blocking, 1.5 mL of each sample of collected infection media was incubated with 15 μL of the human cytokine array detection antibody cocktail at room temperature for 1 hr. After 1 hr, the blocking buffer was poured off and the sample/antibody mixture was incubated with the array membrane at 4 °C overnight. Following incubation with the sample/antibody mixture, the array membrane was washed 3 times with PBS. A streptavidin-HRP conjugate was then added to the membrane and incubated for 30 min on a rocking platform. The array membrane was then washed 3 times with wash buffer to remove any unbound streptavidin-HRP. After the membrane was sufficiently washed it was incubated with a chemiluminescent substrate solution and results were analyzed using the G:Box gel imaging platform.

### ELISA

HCT116 cells were seeded to confluence in 24-well plates (2 x 10^5^ cells/well at 100% confluence) and *F. nucleatum* was added to 500 μL of these wells at a MOI of 50:1. The plates were then incubated at 37 °C, 5% CO_2_ for four hours. The media was then collected from the wells, sterile filtered using a 0.2 μm filter (Millipore Sigma) and diluted to concentrations within the range of the R&D Systems DuoKit ELISA to analyze human IL-8 and CXCL1 concentrations. For ELISAs detecting mouse CCL3 and CXCL2, 1 x 10^6^ fresh neutrophils in suspension were used as described for the HCT116 protocol.

### Transwell HCT116 migration assays

Transwell HCT116 migration assays were performed with Corning 8 μm Transwell inserts in 24-well plates. HCT116 cells were first cultured in McCoy’s 5A (ATCC) to 90% confluence, followed by collection with trypsin Cells were then stained using CellTrackerRed (ThermoFisher) and resuspended at a concentration of 2 x 10^6^ cells/mL in media supplemented with 1% FBS. 100 μL (2 x 10^5^ cells) of the cell suspension was added to the top chamber of the 8 μm Transwell insert pre-coated with 100 μL Matrigel (Corning) (250 μg/mL). The lower chamber contained 600 μL of media with 1% FBS in addition to adding the following: 1) chemokines; purified recombinant human IL-8 (ThermoFisher Scientific) and CXCL1 (Sigma-Aldrich), individually or together at a concentration of 100 ng/mL; 2) conditioned and concentrated media obtained from four hour *F. nucleatum* infections of HCT116 cells. To prepare concentrated media for each sample, three T-75 flasks with confluent HCT116 cells were used. Before infection, the complete media in each flask was replaced with 10mL serum-free and PenStrep-free media. The bacteria resuspended in serum-free media were added at 50:1 MOI, and the flasks were incubated in a hypoxic chamber (1% O_2_) for four hours. The media containing the cell secretions was collected and pooled from three flasks, spun down at 3000xG for five minutes and passed through a 0.22μm filter. The samples were then concentrated from 30 mL to 1.5 mL using a 3000 MWCO Amplico concentrator (Millipore Sigma) at 4 °C. The resulting sample was used for ELISA and Transwell migration assays. The Transwells were incubated at 37 °C, 5% CO_2_ for 6-16 hours, after which the cells on the top were removed using a cotton-tipped applicator and migrated cells at the bottom of the membrane were stained with DAPI and imaged on a Zeiss LSM 800 confocal microscope using a 10X objective. ImageJ was used to count the number of cells in 5 representative images per Transwell in biological triplicate (*69*).

### Depletion of IL-8 and CXCL1 from conditioned media

Conditioned and concentrated media was obtained as described. Human IL-8/CXCL8 biotinylated antibody (R&D Systems BAF208) and Human/Primate CXCL1/GROα/KC/CINC-1 biotinylated antibody (R&D Systems BAF275) were added to the media to a final concentration of 40 ng/mL and incubated at room temperature with gentle shaking for 30 minutes. 150 μL of magnetic streptavidin particles (Sigma-Aldrich 11641778001) were first washed twice with PBS and spun down at 1500 xG and added to the solution. The mixture was incubated at room temperature with gentle shaking for another 30 minutes. The samples were then spun down at 1500 xG, and the supernatant was collected containing the conditioned media with reduced cytokines. An ELISA quantified and confirmed the depletion. This media was subsequently used in transwell migration assays as described.

### Statistical analysis

All statistical analysis was performed in GraphPad Prism Version 8.2.1. For single analysis, an unpaired Student’s *t* test was used. For grouped analyses, Two-way ANOVA was used. In each case, the following *P* values correspond to star symbols in figures: ^ns^*P* >0.05, \**P* < 0.05, \*\**P* < 0.01, *** *P* < 0.001, ****P < 0.0001. To obtain statistics, all studies were performed as three independent biological experiments.

## Author contributions

*Michael A. Casasanta*, Data curation, Methodology, Formal Analysis, Writing-review and editing; *Christopher C. Yoo*) Data curation, Methodology, Formal Analysis, Writing-review and editing; *Barath Udayasuryan_t_* Data curation, Methodology, Formal Analysis, Writing-review and editing; *Blake E. Sanders*) Data curation, Writing-review and editing; *Ariana Umaña_t_* Data curation, Writing-review and editing; *Yao Zhang*) Data curation, Writing-review and editing; *Huaiyao Peng_t_* Data curation, Writing-review and editing; *A. Jane Duncan*, Data curation, Writing-review and editing; *Yueying Wang*) Data curation, Writing-review and editing; *Liwu Li,* Conceptualization, Formal analysis, Supervision, Funding acquisition, Validation, Methodology, Writing-review and editing; *Scott S. Verbridge*) Conceptualization, Formal analysis, Supervision, Funding acquisition, Validation, Methodology, Project administration, review and editing; *Daniel J. Slade*) Conceptualization, Data curation, Formal analysis, Supervision, Funding acquisition, Validation, Methodology, Project administration, Writing-original draft, review and editing.

## Data and materials availability

Materials are available upon reasonable request or in the future through the Addgene plasmid repository. All data needed to evaluate conclusions are presented in the paper or Supplementary Materials, but additional raw data can be accessed on our Open Science Framework repository at: https://osf.io/kbj2h/

## SUPPLEMENTARY MATERIALS

**Fig. S1.**
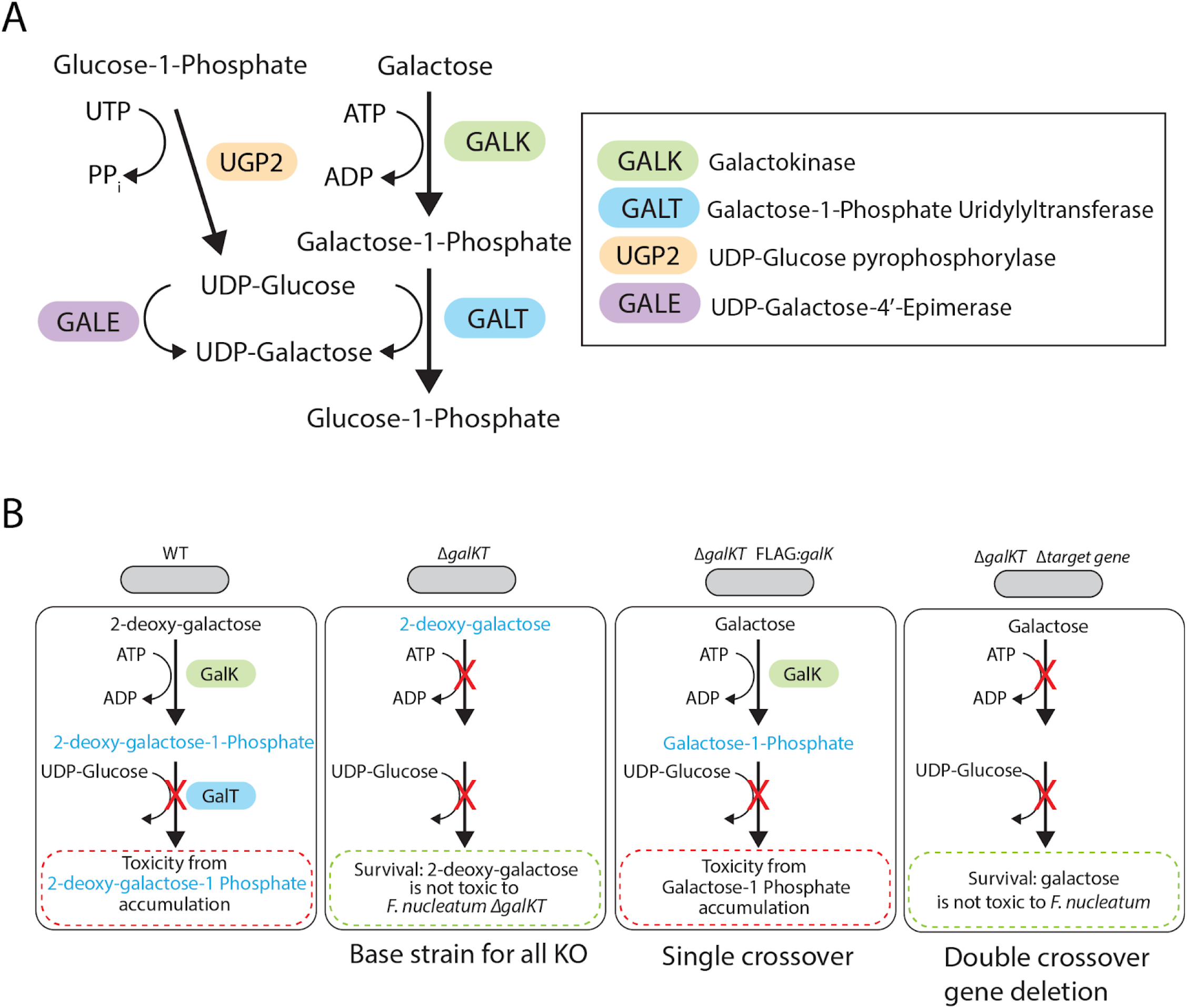
Utilizing the Leloir pathway for targeted gene deletions in bacteria. (**A**) The Leloir pathway of galactose catabolism. (**B**) Graphical representation of the different selection stages of *F. nucleatum* gene deletions and how the GalK and GalT affect cellular survival in the presence of galactose and 2-deoxy-galactose.

**Fig. S2.**
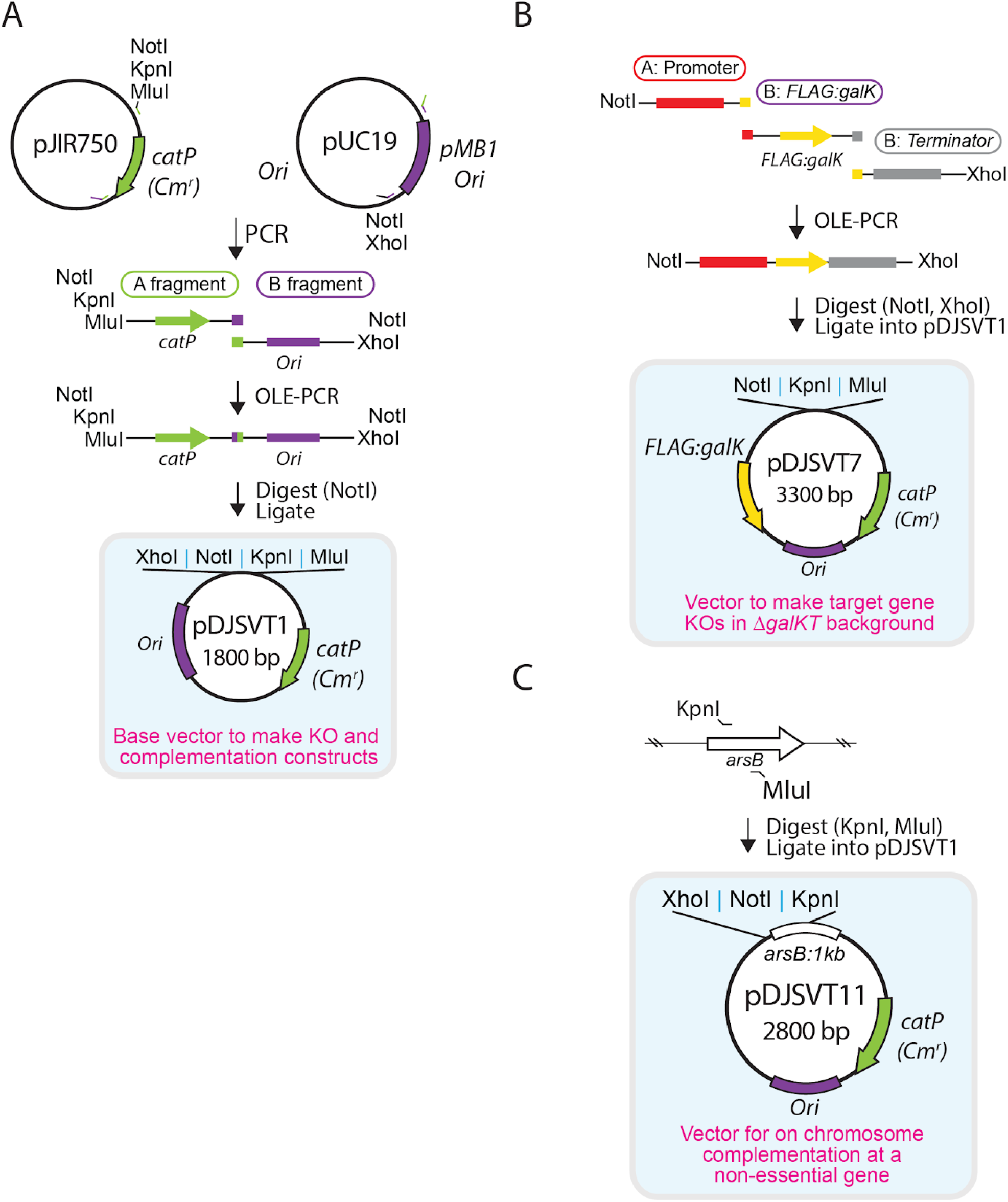
Development of vector for markerless gene deletion in *Fusobacterium nucleatum*. (**A**) Method to create pDJSVT1; the base vector for all gene deletion constructs that consists of an *E. coli* origin and chloramphenicol/thiamphenicol resistance and a GC rich multiple cloning site for efficient cloning of AT rich *F. nucleatum* DNA. (**B**) pDJSVT7 plasmid incorporation of a constitutively active *FLAG:galK* gene for selection on 2-deoxy-galactose. (**C**) Development of a chromosomal complementation vector that incorporates the entire pDJSVT11 plasmid onto the chromosome at the *arsB* gene in *F. nucleatum* 23726.

**Fig. S3.**
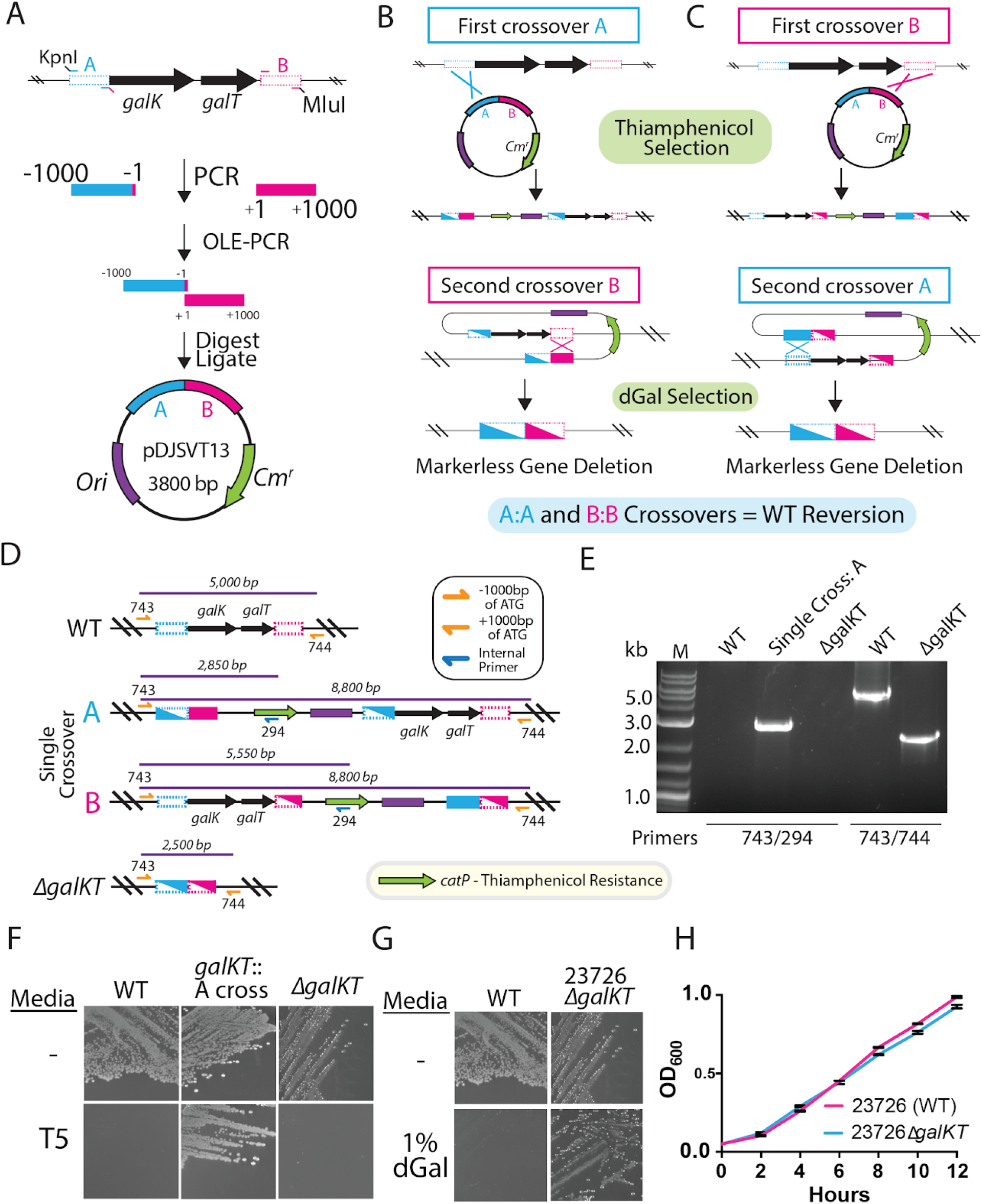
Deletion of the *galKT* gene operon in *F. nucleatum* 23726. (**A**) Development of pDJSVT13, the plasmid for *galKT* gene deletion, by inserting 1000 bp *galKT* flanking sequences into pDJSVT1. (**B**) Representation of single-crossover homologous recombination onto the *F. nucleatum* 23726 chromosome with either the A (upstream) or (**C**) B (downstream) 1000 bp DNA fragments in pDJSVT13 followed by showing how the double crossover results in complete plasmid excision. (**D**) Map of chromosomal plasmid incorporation after the initial single-crossover insertion. (**E**) PCR verification of single-cross pDJSVT13 and subsequent double-crossover deletion of the *galKT* operon. (**F**) Plating on 5 μg/mL thiamphenicol (T5) (single-crossover selection) and (**G**) 1% 2-deoxy-galactose (double-crossover), resulting in galKT operon deletion. (**H**) Validation that *F. nucleatum* 23726 Δ*galKT* grows the same as WT *F. nucleatum* 23726.

**Fig. S4.**
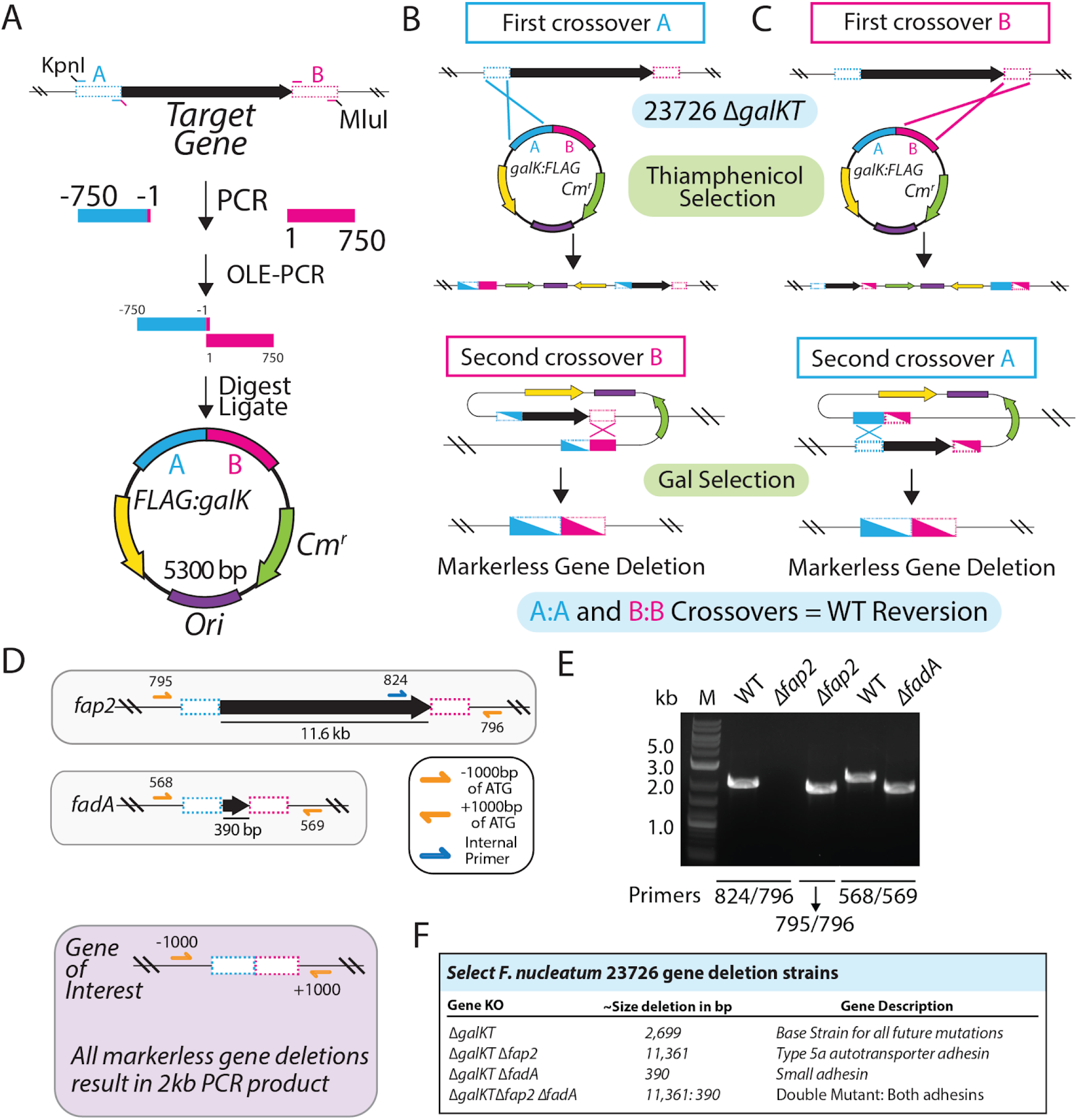
Target gene deletion in *F. nucleatum* 23726 Δ*galKT.* (**A**) Development of all target gene deletion plasmids (Table S2) by inserting 750 bp from target gene flanking sequences into pDJSVT7. (**B**) Representation of single-crossover homologous recombination onto the *F. nucleatum* 23726 chromosome with either the A (upstream) or (**C**) B (downstream) 750 bp DNA fragments. Double crossover results are shown during complete plasmid excision. (**D**) *fap2* and *fadA* as examples of gene excision. KO validation primers are shown flanking genes at −1000 and +1000 bp. This results in 2kb PCR products as shown in (**E**) for the *fap2* and *fadA* gene deletions. (**F**) Table of select deleted genes in this study and their function.

**Fig. S5.**
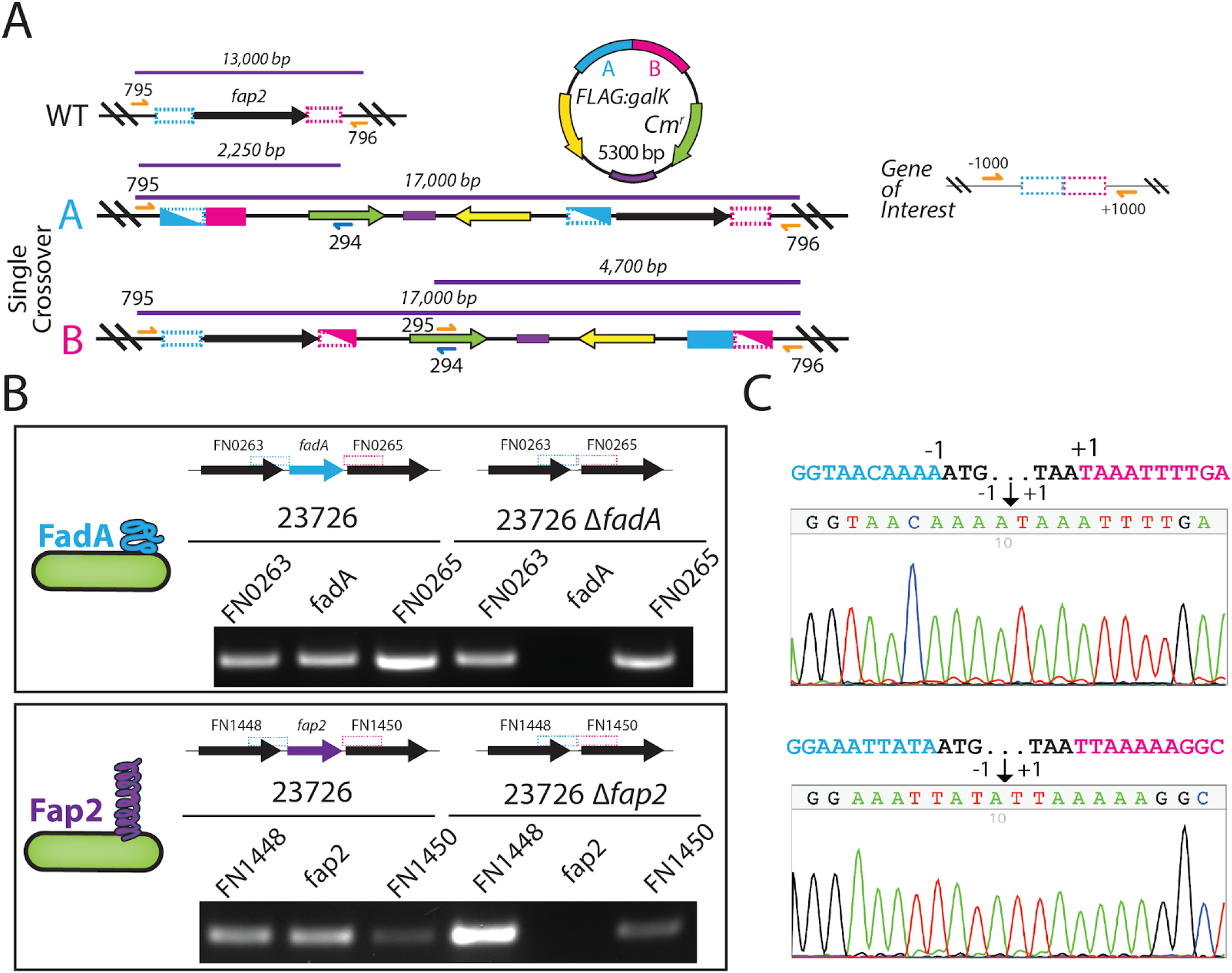
Validation of markerless *fap2* and *fadA* gene deletions in *F. nucleatum* 23726 Δ*galKT.* (**A**) Map of chromosomal plasmid incorporation after the initial single-crossover insertion for *fap2*. (**B**) RT-PCR of *fadA* and *fap2* with their two surrounding genes showing loss of gene transcription due to gene deletion. (**C**) DNA sequencing with primers sitting −250 and +250 from the start codon of *fadA* and *fap2*. All constructs created are 100% accurate and show perfect excision and recission of the genome to the −1 and +1 base for each gene.

**Fig. S6.**
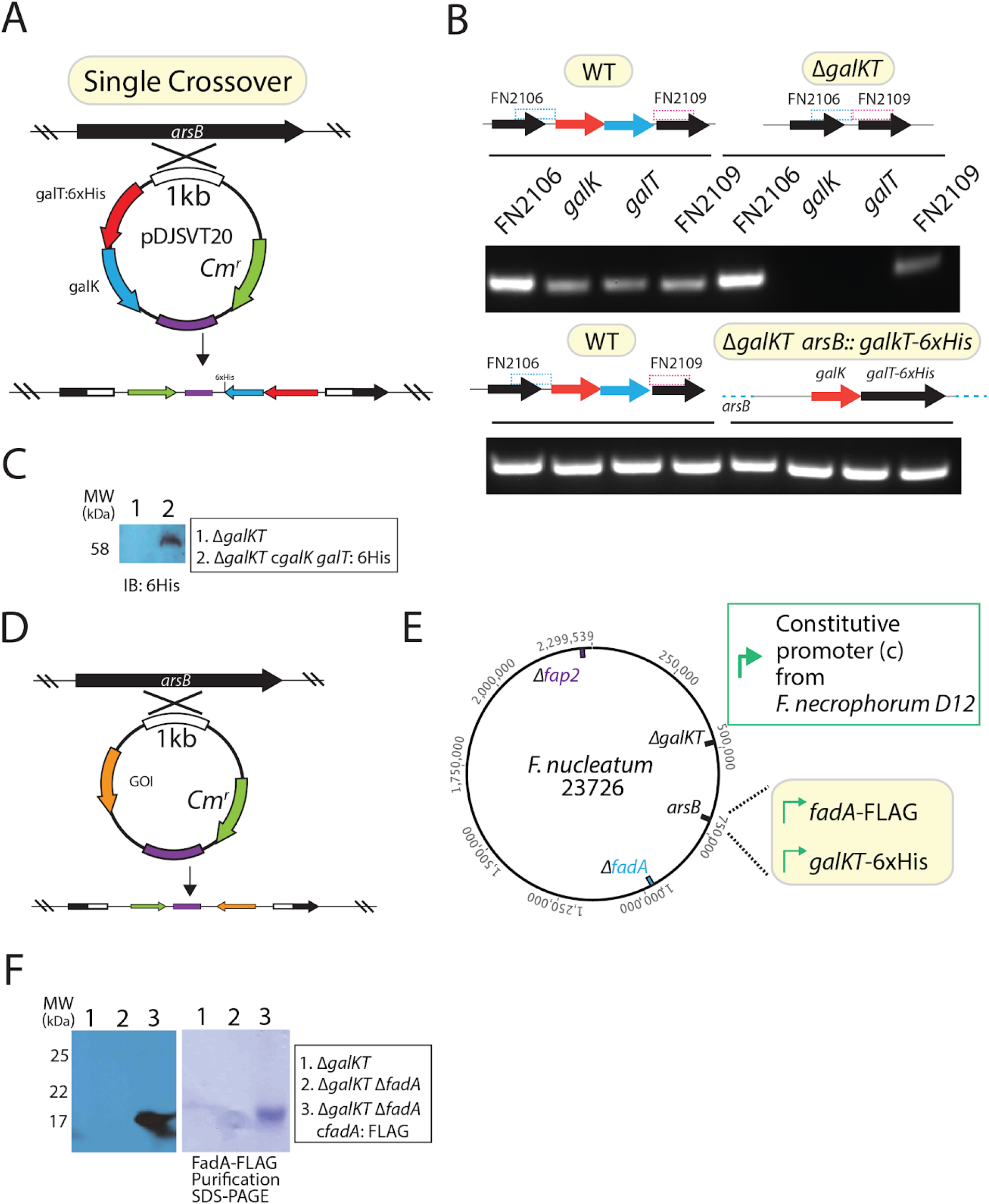
Complementation of gene deletions at the static *arsB* gene site in *F. nucleatum* 23726. (**A**) pDJSVT20 plasmid created for complementing Δ*galKT* with galKT::6xHis. (**B**) RT-PCR of Δ*galKT* and Δ*galKT galKT::6xHis* and the two genes flanking the operon (FN2106, FN2109). (**C**) Western blot verification of *galKT::6sHis* complementation and constitutive expression of GalT::6xHis. (**D**) pDJSVT11 base vector depicted with target gene of interest (GOI) complementation and how it incorporates into the chromosome at *arsB*. (**E**) Potential sites for complementation of gene deletions used in this study. However, we did not complement the Δ*fap2* strain due to the extreme difficulties of complementing large genes (12kb). (**F**) Complementation of Δ*fadA* with *fadA:*: FLAG and western blot verification of constitutive expression of FadA::FLAG.

**Fig. S7.**
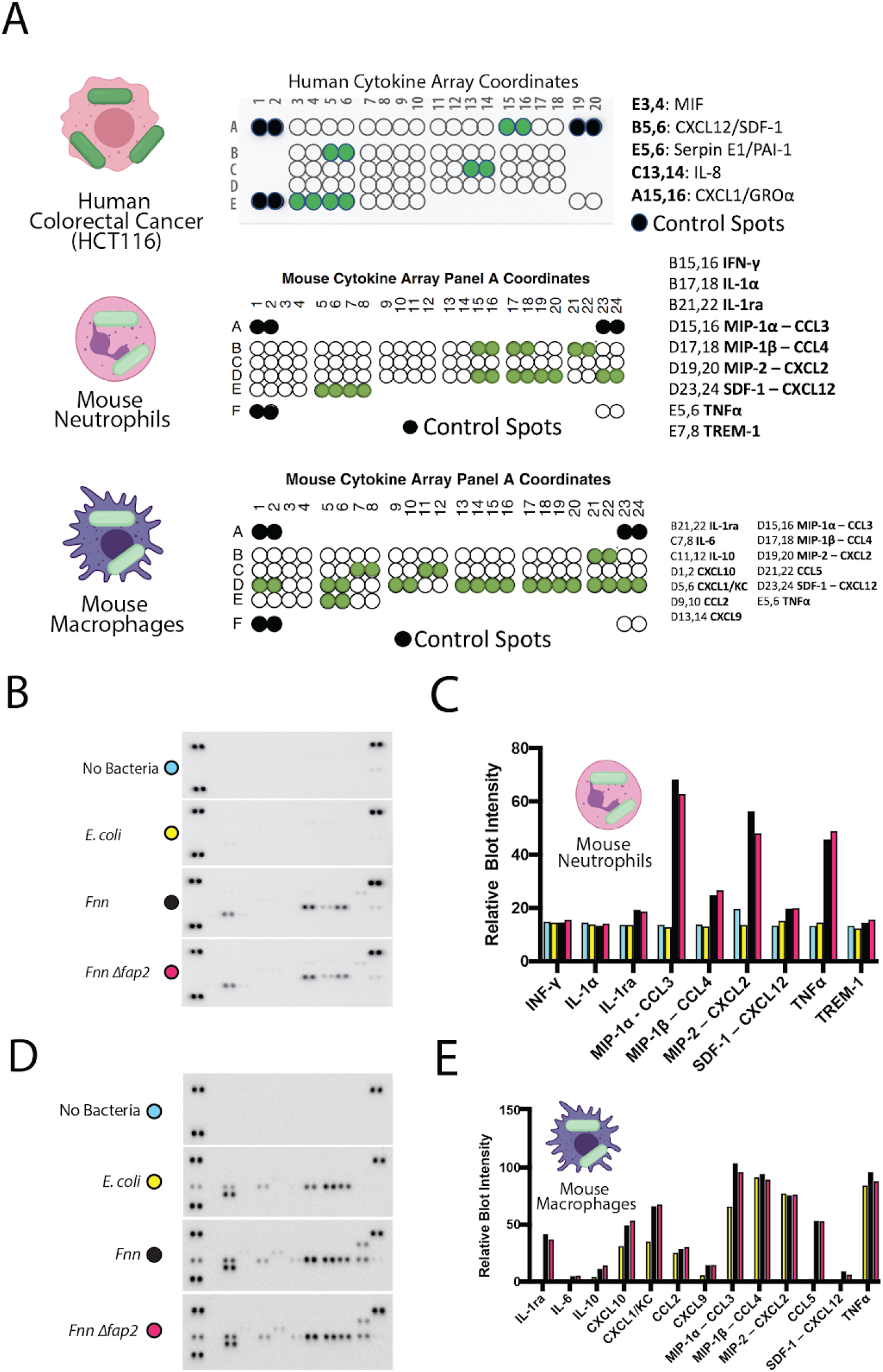
Human and mouse cytokine arrays to detect *F. nucleatum* induced immune signaling. (**A**) Cytokine array coordinates for human and mouse array with green dots representing spots analyzed due to detected expression. Gene names are shown to the right, and full coordinates for all spots can be found at R&D Systems (R&D: Proteome Profiler Human Cytokine Array (ARY005B), Proteome Profiler Mouse Cytokine Array Kit, Panel A (ARY006)). (**B**) Raw blots for mouse cytokines secreted by mouse neutrophils after infection with *F. nucleatum* 23726 at 50:1 MOI for four hours. (**C**) Quantitation of relative spot intensity compared to the control blot of uninfected neutrophils. (**D**) Raw blots for mouse cytokines secreted by mouse macrophages after infection with *F. nucleatum* 23726 at 50:1 MOI for four hours. (**E**) Quantitation of relative spot intensity compared to the control blot of uninfected macrophages.

**Supplementary Table 1:**
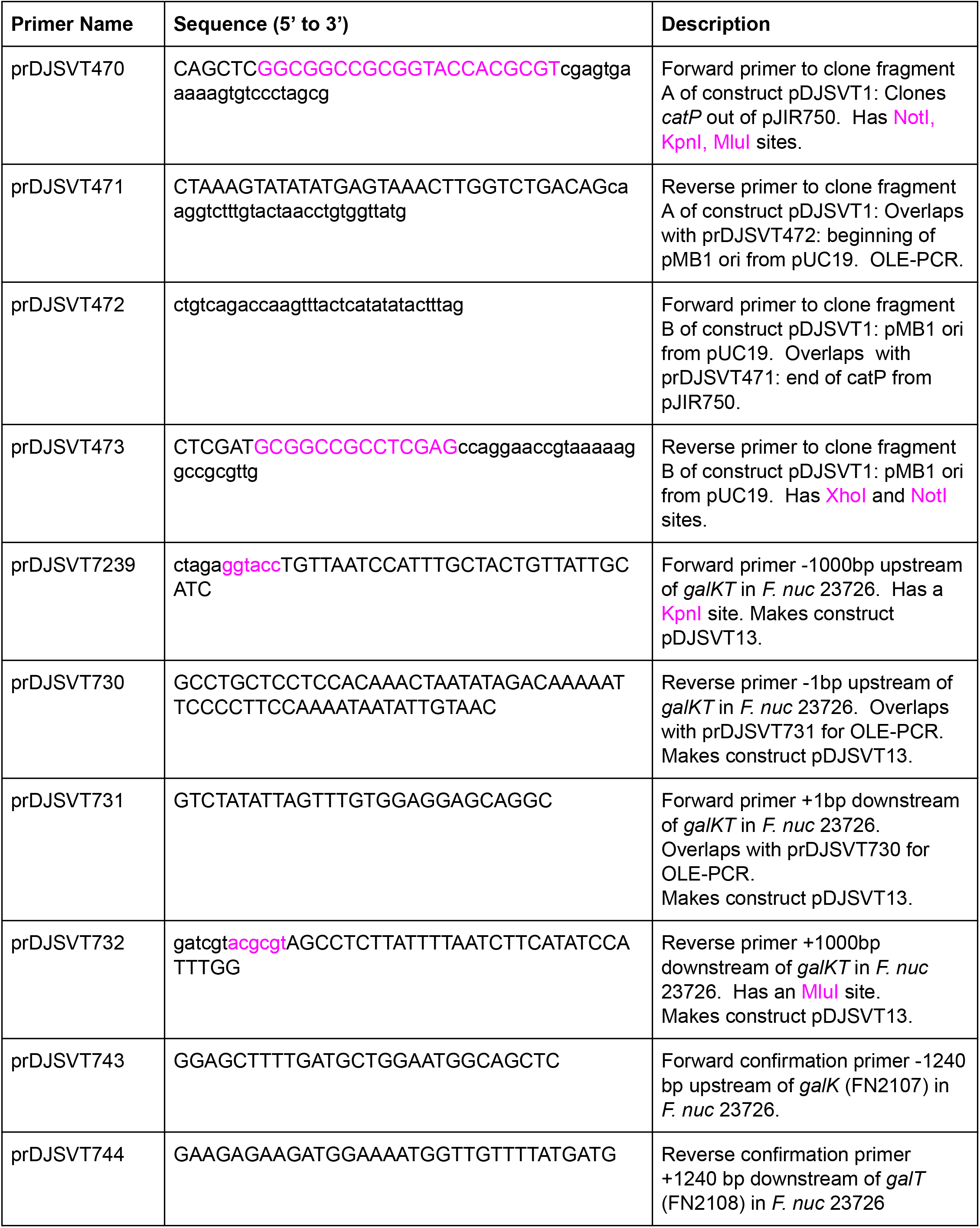
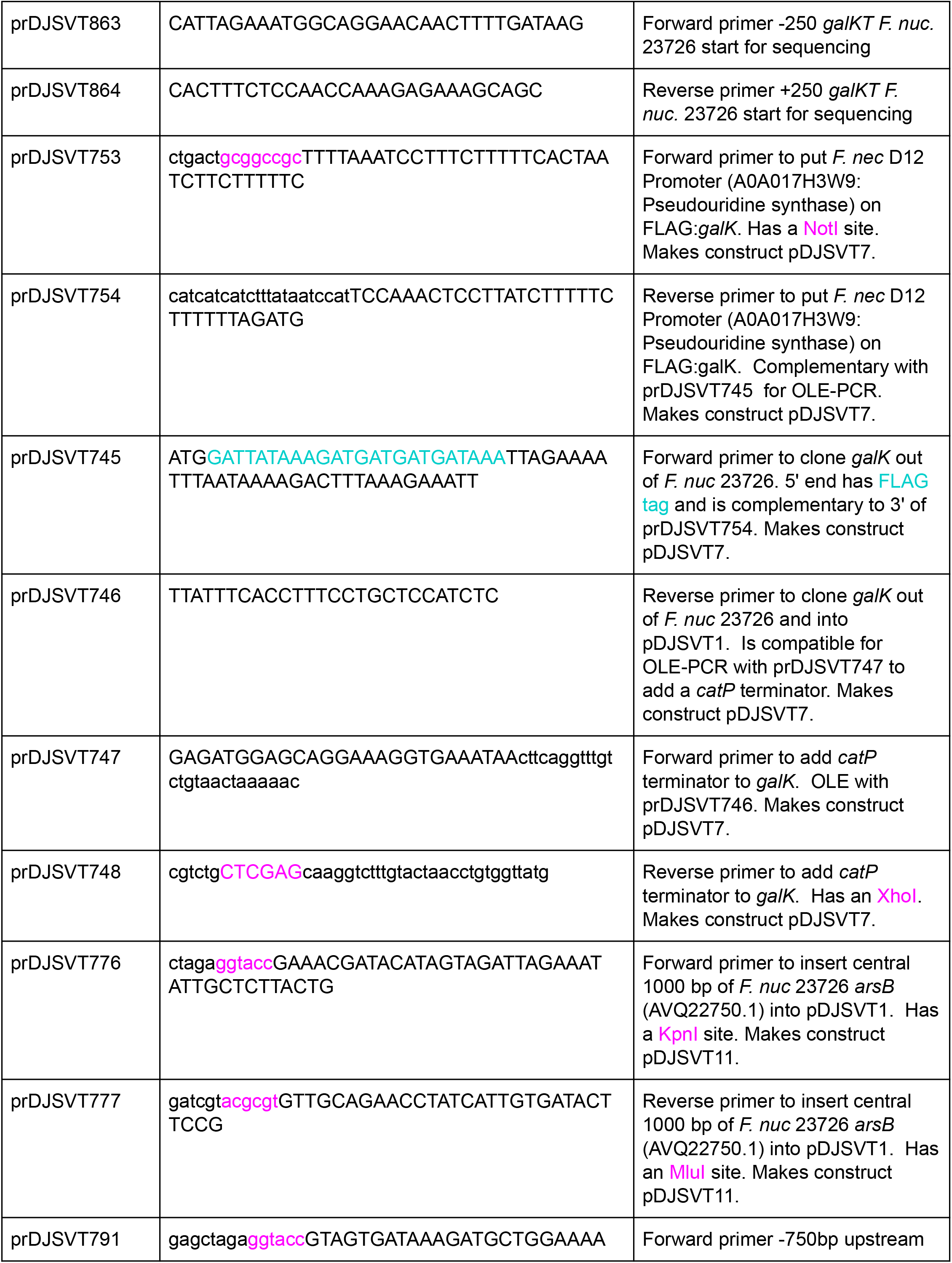
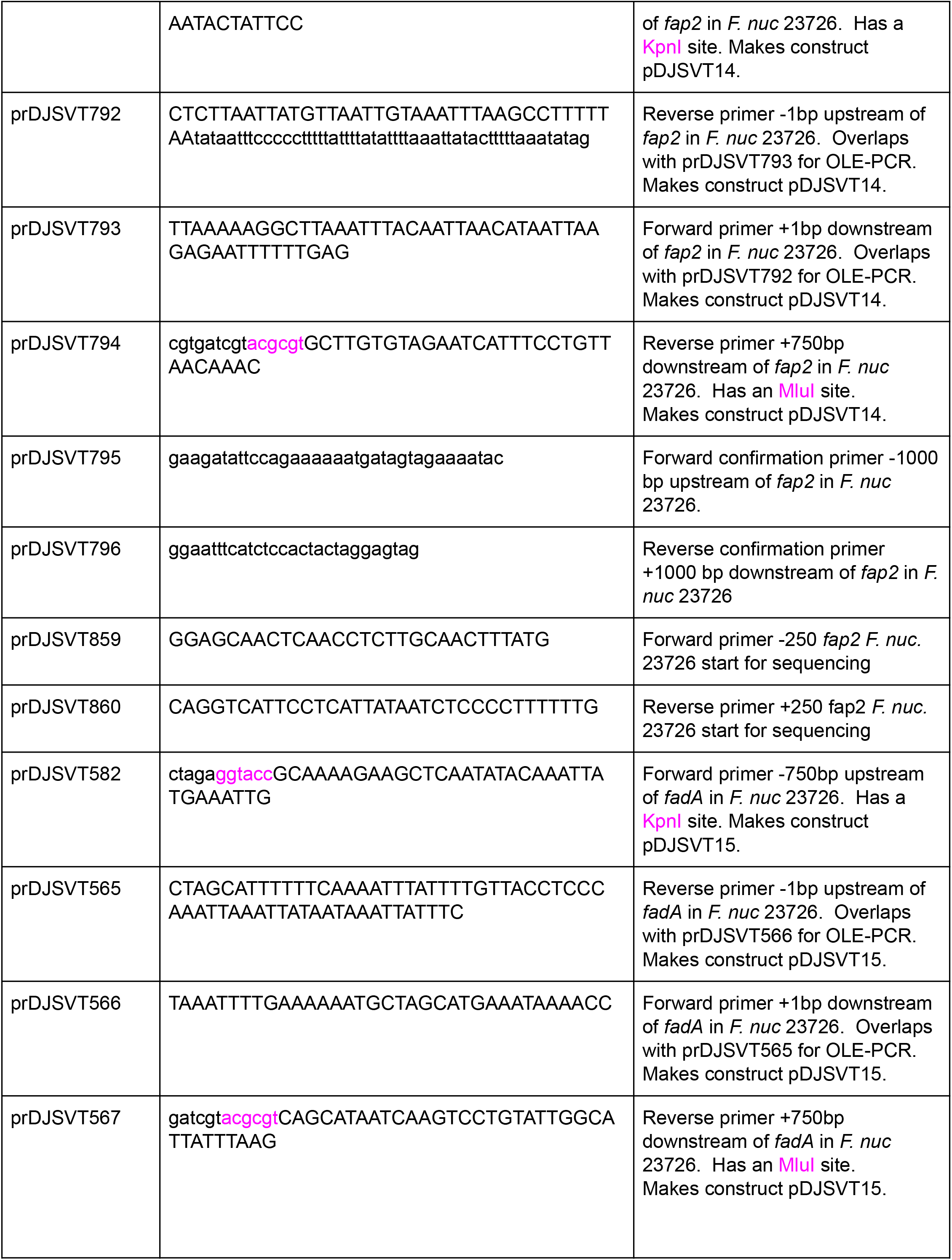
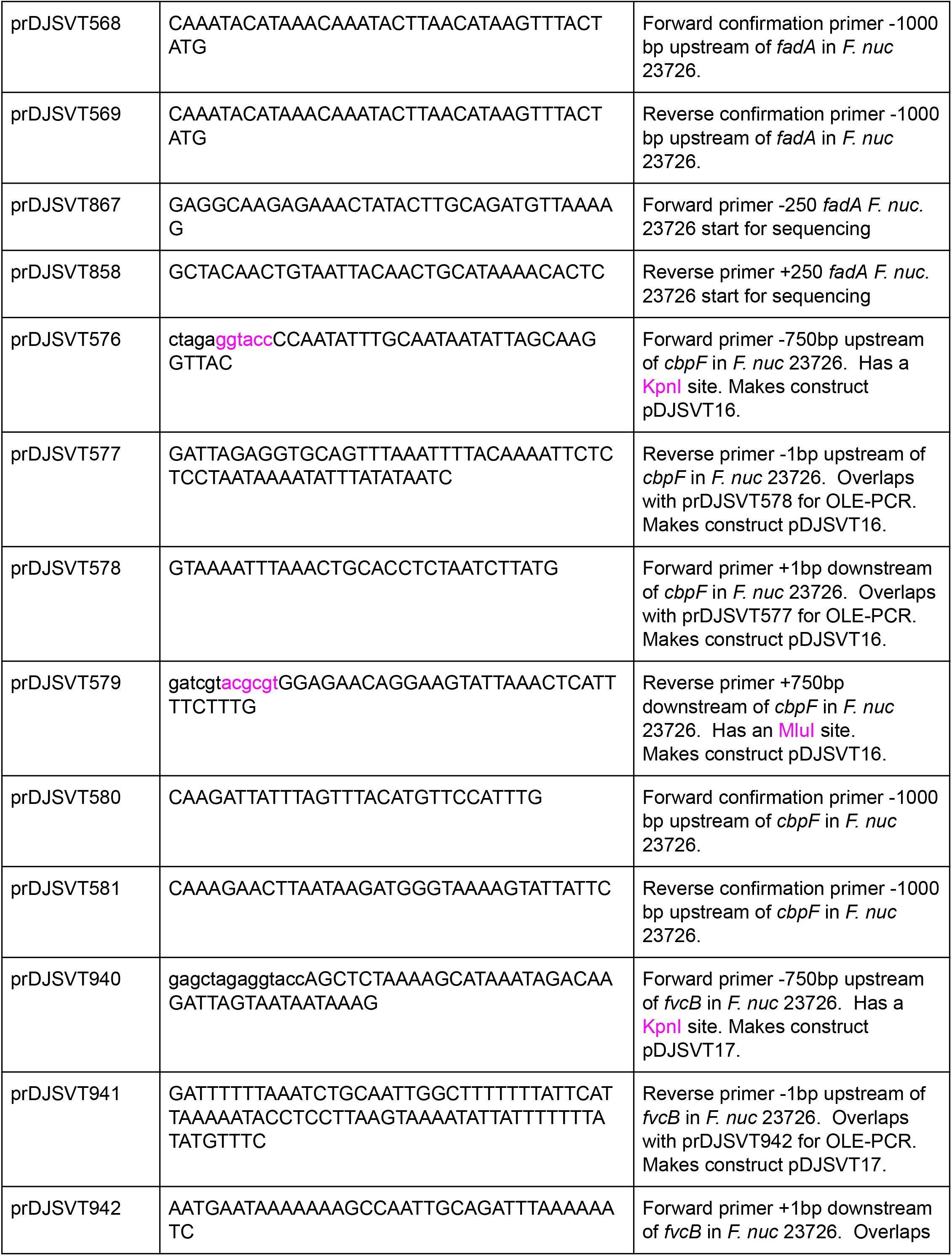
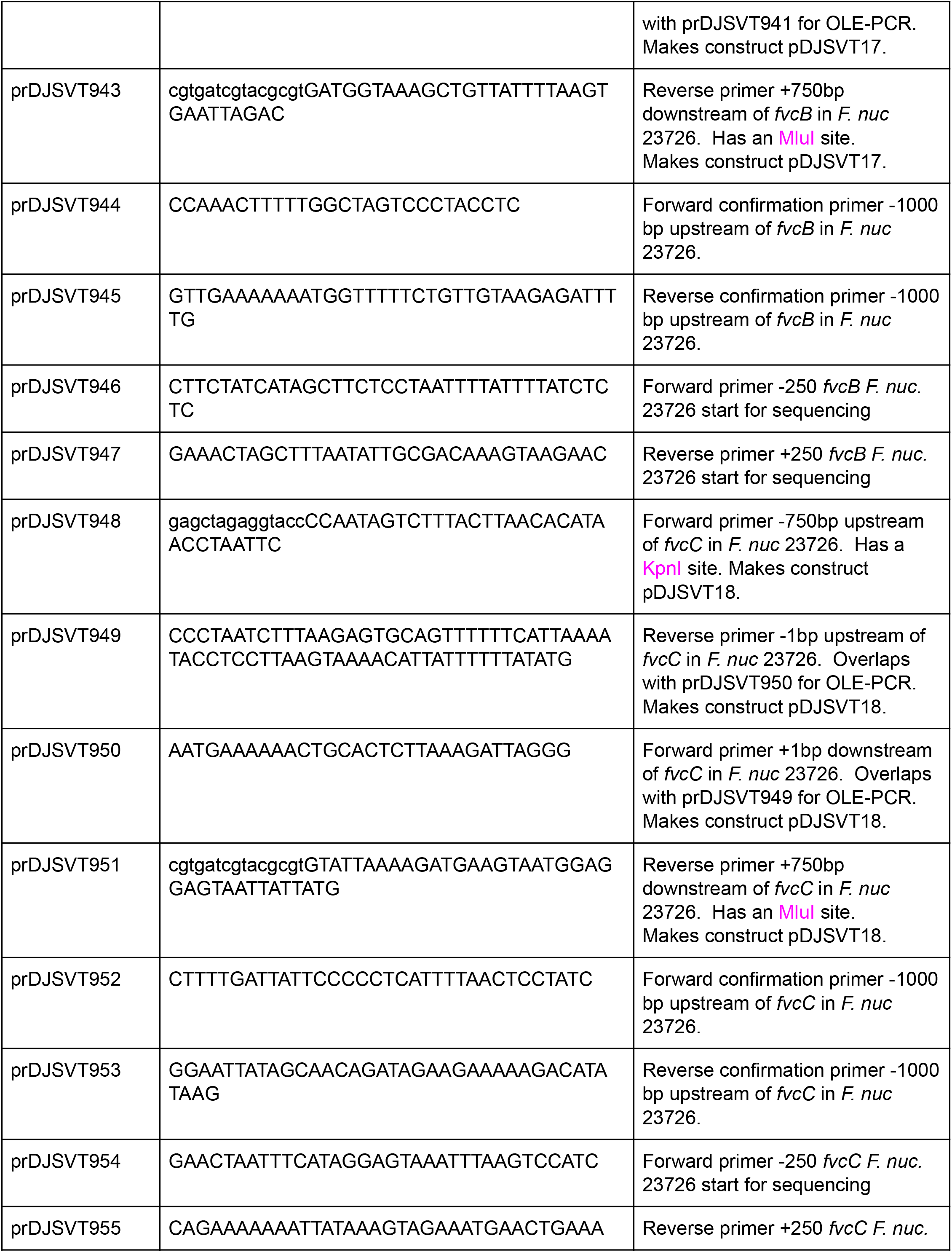
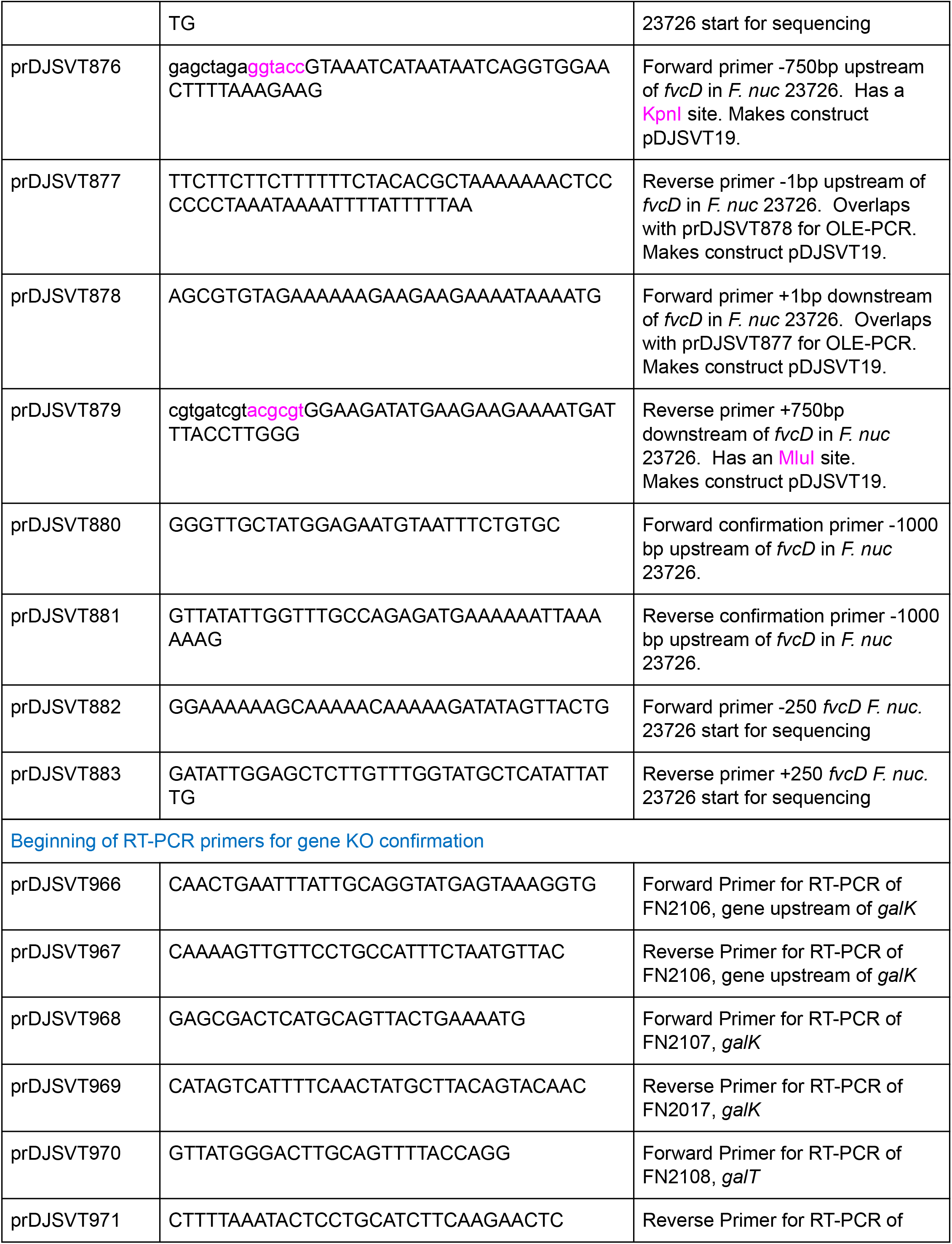
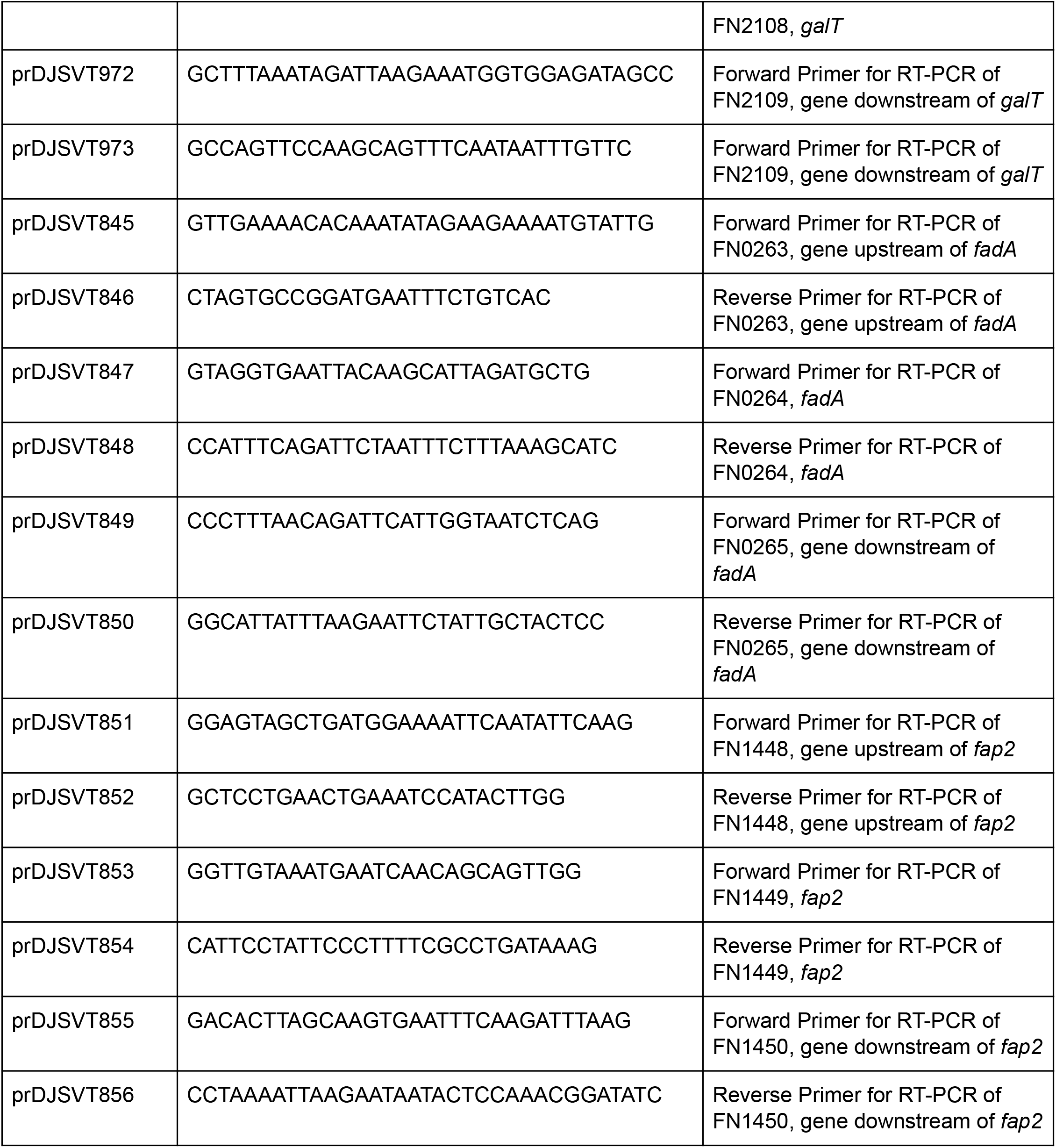
Primers used in this study.

**Supplementary Table 2:**
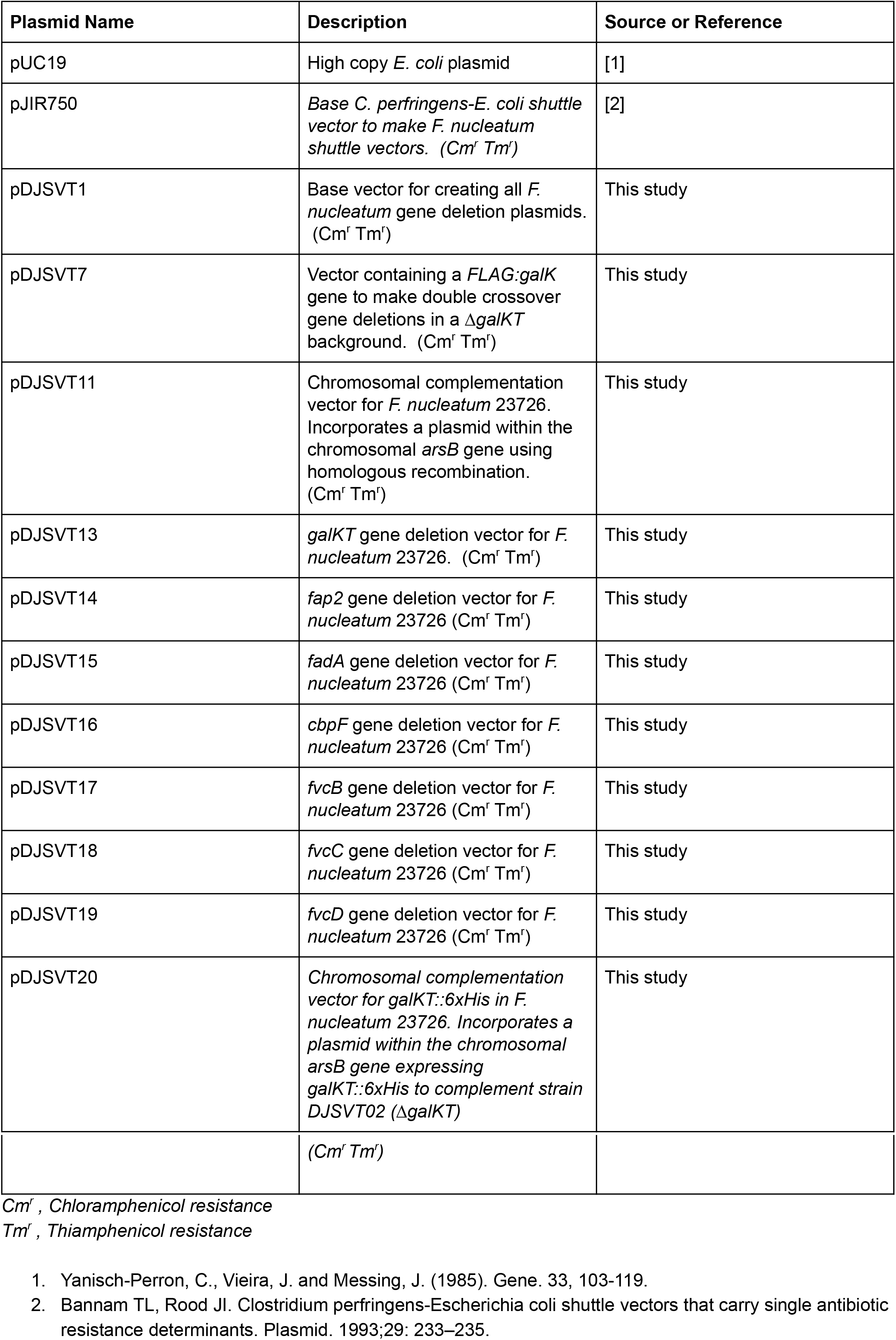
Plasmids used in this study.

**Supplementary Table 3:**
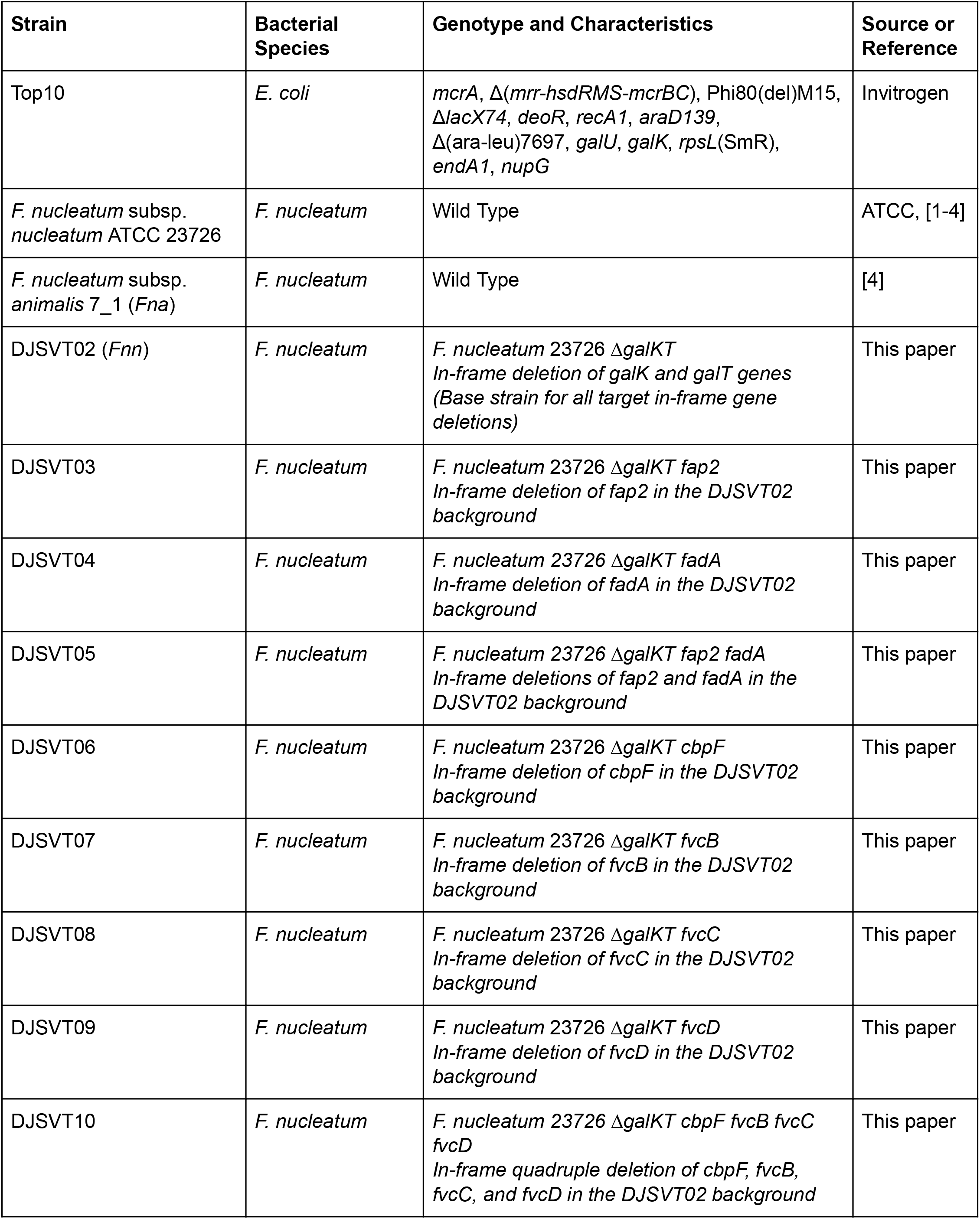
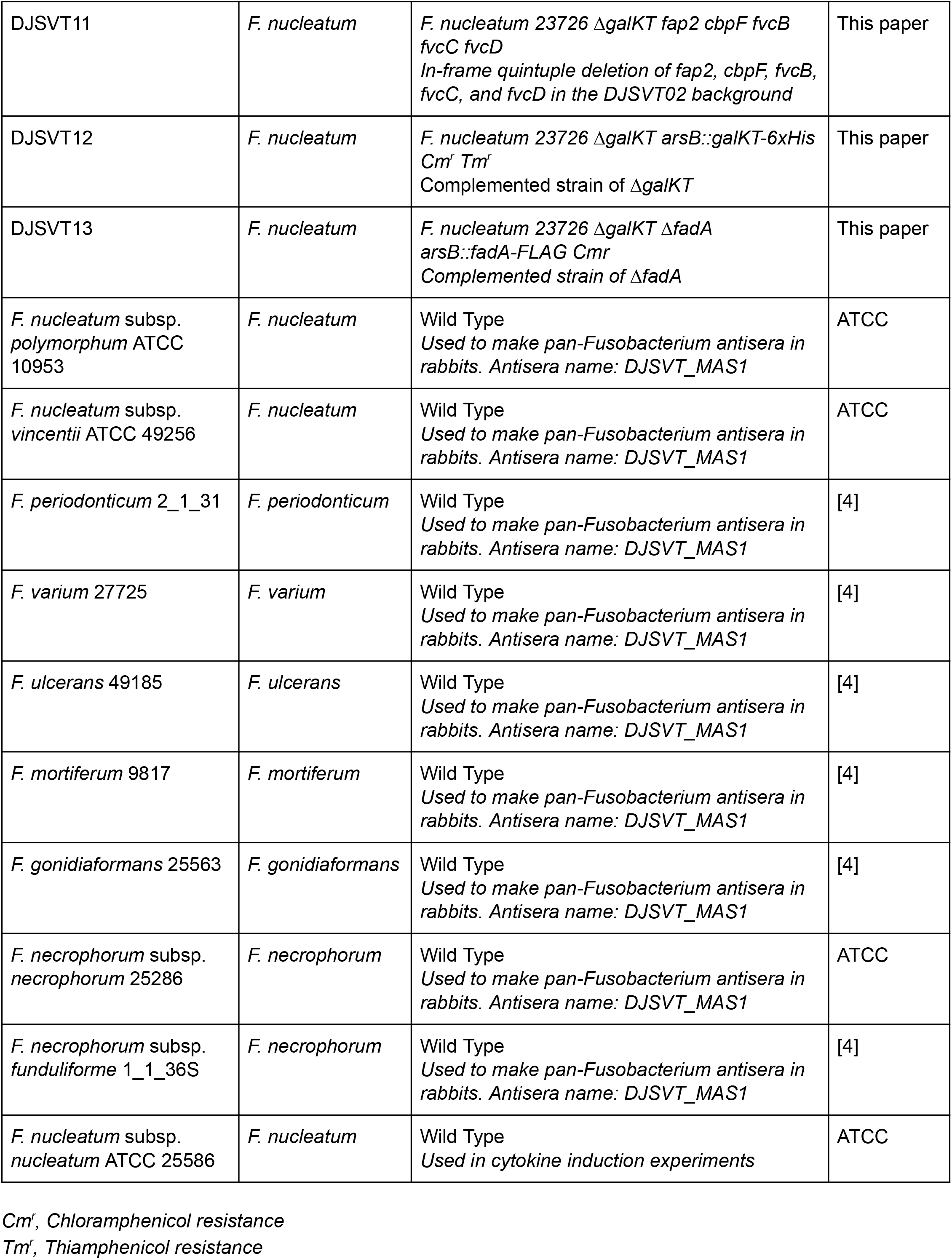
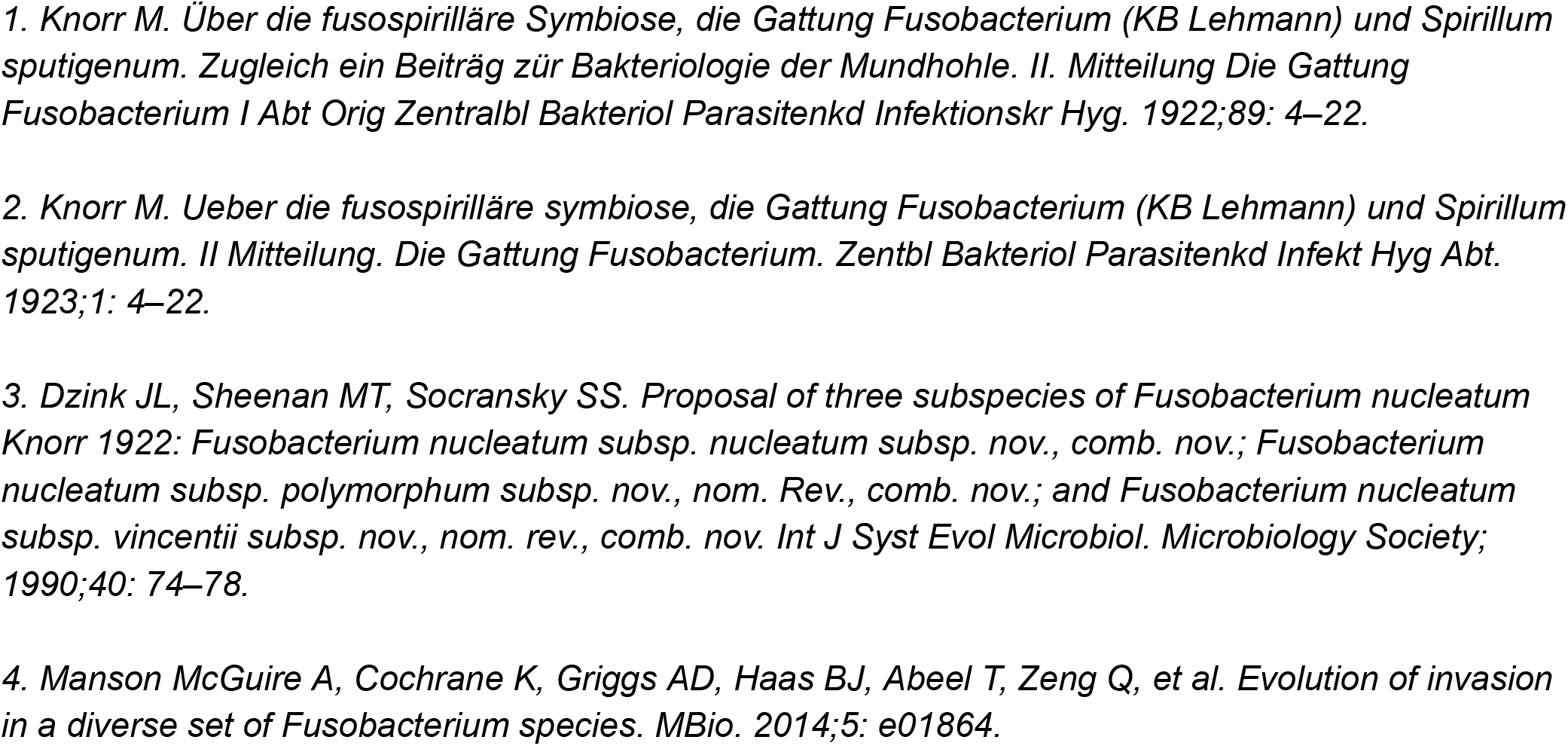
Bacterial strains used in this study.

## REFERENCES

1. D. van Elsland, J. Neefjes, Bacterial infections and cancer. EMBO Rep. 19 (2018), doi:10.15252/embr.201846632.

2. A. Gagnaire, B. Nadel, D. Raoult, J. Neefjes, J.-P. Gorvel, Collateral damage: insights into bacterial mechanisms that predispose host cells to cancer. Nat. Rev. Microbiol. 15, 109–128 (2017).

3. H. L. Kaufman, F. J. Kohlhapp, A. Zloza, Oncolytic viruses: a new class of immunotherapy drugs. Nat. Rev. Drug Discov 14, 642–662 (2015).

4. C. A. Brennan, W. S. Garrett, Fusobacterium nucleatum - symbiont, opportunist and oncobacterium. Nat. Rev. Microbiol. (2018), doi:10.1038/s41579-018-0129-6.

5. Y. W. Han, Fusobacterium nucleatum: a commensal-turned pathogen. Curr. Opin. Microbiol. 23, 141–147 (2015).

6. M. R. Rubinstein, X. Wang, W. Liu, Y. Hao, G. Cai, Y. W. Han, Fusobacterium nucleatum promotes colorectal carcinogenesis by modulating E-cadherin/β-catenin signaling via its FadA adhesin. Cell Host Microbe, 14, 195–206 (2013).

7. J. Abed, J. E. M. Emgård, G. Zamir, M. Faroja, G. Almogy, A. Grenov, A. Sol, R. Naor, E. Pikarsky, K. A. Atlan, A. Mellul, S. Chaushu, A. L. Manson, A. M. Earl, N. Ou, C. A. Brennan, W. S. Garrett, G. Bachrach, Fap2 Mediates Fusobacterium nucleatum Colorectal Adenocarcinoma Enrichment by Binding to Tumor-Expressed Gal-GalNAc. Cell Host Microbe. 20, 215–225 (2016).

8. M. Castellarin, R. L. Warren, J. D. Freeman, L. Dreolini, M. Krzywinski, J. Strauss, R. Barnes, P. Watson, E. Allen-Vercoe, R. A. Moore, R. A. Holt, Fusobacterium nucleatum infection is prevalent in human colorectal carcinoma. Genome Res. 22, 299–306 (2012).

9. A. D. Kostic, D. Gevers, C. S. Pedamallu, M. Michaud, F. Duke, A. M. Earl, A. I. Ojesina, J. Jung, A. J. Bass, J. Tabernero, J. Baselga, C. Liu, R. A. Shivdasani, S. Ogino, B. W. Birren, C. Huttenhower, W. S. Garrett, M. Meyerson, Genomic analysis identifies association of Fusobacterium with colorectal carcinoma. Genome Res. 22, 292–298 (2012).

10. S. Bullman, C. S. Pedamallu, E. Sicinska, T. E. Clancy, X. Zhang, D. Cai, D. Neuberg, K. Huang, F. Guevara, T. Nelson, O. Chipashvili, T. Hagan, M. Walker, A. Ramachandran, B. Diosdado, G. Serna, N. Mulet, S. Landolfi, S. Ramon Y Cajal, R. Fasani, A. J. Aguirre, K. Ng, E. Élez, S. Ogino, J. Tabernero, C. S. Fuchs, W. C. Hahn, P. Nuciforo, M. Meyerson, Analysis of Fusobacterium persistence and antibiotic response in colorectal cancer. Science, 358, 1443–1448 (2017).

11. S. Chen, T. Su, Y. Zhang, A. Lee, J. He, Q. Ge, L. Wang, J. Si, W. Zhuo, L. Wang, Fusobacterium nucleatum promotes colorectal cancer metastasis by modulating KRT7-AS/KRT7. Gut Microbes, 1–15 (2020).

12. Y. Chen, Y. Chen, J. Zhang, P. Cao, W. Su, Y. Deng, N. Zhan, X. Fu, Y. Huang, W. Dong, Fusobacterium nucleatum Promotes Metastasis in Colorectal Cancer by Activating Autophagy Signaling via the Upregulation of CARD3 Expression. Theranostics, 10, 323–339 (2020).

13. H. Verbeke, S. Struyf, G. Laureys, J. Van Damme, The expression and role of CXC chemokines in colorectal cancer. Cytokine Growth Factor Rev. 22, 345–358 (2011).

14. C. Zhuo, X. Wu, J. Li, D. Hu, J. Jian, C. Chen, X. Zheng, C. Yang, Chemokine (C-X-C motif) ligand 1 is associated with tumor progression and poor prognosis in patients with colorectal cancer. Biosci. Rep. 38 (2018), doi:10.1042/BSR20180580.

15. A. Li, M. L. Varney, R. K. Singh, Constitutive expression of growth regulated oncogene (gro) in human colon carcinoma cells with different metastatic potential and its role in regulating their metastatic phenotype. Clin. Exp. Metastasis. 21, 571–579 (2004).

16. A. Li, M. L. Varney, J. Valasek, M. Godfrey, B. J. Dave, R. K. Singh, Autocrine role of interleukin-8 in induction of endothelial cell proliferation, survival, migration and MMP-2 production and angiogenesis. Angiogenesis? 8, 63–71 (2005).

17. H. Ogata, A. Sekikawa, H. Yamagishi, K. Ichikawa, S. Tomita, J. Imura, Y. Ito, M. Fujita, M. Tsubaki, H. Kato, T. Fujimori, H. Fukui, GROα promotes invasion of colorectal cancer cells. Oncol. Rep. 24, 1479–1486 (2010).

18. D. Wang, H. Sun, J. Wei, B. Cen, R. N. DuBois, CXCL1 Is Critical for Premetastatic Niche Formation and Metastasis in Colorectal Cancer. Cancer Res. 77, 3655–3665 (2017).

19. C. Rubie, V. O. Frick, S. Pfeil, M. Wagner, O. Kollmar, B. Kopp, S. Graber, B. M. Rau, M. K. Schilling, Correlation of IL-8 with induction, progression and metastatic potential of colorectal cancer. World J. Gastroenterol. 13, 4996–5002 (2007).

20. L.-C. Chen, C.-Y. Hao, Y. S. Y. Chiu, P. Wong, J. S. Melnick, M. Brotman, J. Moretto, F. Mendes, A. P. Smith, J. L. Bennington, D. Moore, N. M. Lee, Alteration of gene expression in normal-appearing colon mucosa of APC(min) mice and human cancer patients. Cancer Res. 64, 3694–3700 (2004).

21. Y. S. Lee, I. Choi, Y. Ning, N. Y. Kim, V. Khatchadourian, D. Yang, H. K. Chung, D. Choi, M. J. LaBonte, R. D. Ladner, K. C. Nagulapalli Venkata, D. O. Rosenberg, N. A. Petasis, H.-J. Lenz Y. K. Hong, Interleukin-8 and its receptor CXCR2 in the tumour microenvironment promote colon cancer growth, progression and metastasis. Br. J. Cancer 106, 1833–1841 (2012).

22. Q. Liu, A. Li, Y. Tian, J. D. Wu, Y. Liu, T. Li, Y. Chen, X. Han, K. Wu, The CXCL8-CXCR1/2 pathways in cancer. Cytokine Growth Factor Rev. 31, 61–71 (2016).

23. A. Umaña, B. E. Sanders, C. C. Yoo, M. A. Casasanta, B. Udayasuryan, S. S. Verbridge, D. J. Slade, Utilizing Whole Fusobacterium Genomes To Identify, Correct, and Characterize Potential Virulence Protein Families. J. Bacteriol. 201 (2019), doi:10.1128/JB.00273-19.

24. M. Desvaux, A. Khan, S. A. Beatson, A. Scott-Tucker, I. R. Henderson, Protein secretion systems in Fusobacterium nucleatum: genomic identification of Type 4 piliation and complete Type V pathways brings new insight into mechanisms of pathogenesis. Biochim. Biophys. Acta. 1713, 92–112 (2005).

25. N. Dautin, H. D. Bernstein, Protein secretion in gram-negative bacteria via the autotransporter pathway. Annu. Rev. Microbiol. 61, 89–112 (2007).

26. C. Gur, Y. Ibrahim, B. Isaacson, R. Yamin, J. Abed, M. Gamliel, J. Enk, Y. Bar-On, N. Stanietsky-Kaynan, S. Coppenhagen-Glazer, N. Shussman, G. Almogy, A. Cuapio, E. Hofer, D. Mevorach, A. Tabib, R. Ortenberg, G. Markel, K. Miklić, S. Jonjic, C. A. Brennan, W. S. Garrett, G. Bachrach, O. Mandelboim, Binding of the Fap2 protein of Fusobacterium nucleatum to human inhibitory receptor TIGIT protects tumors from immune cell attack. Immunity. 42, 344–355 (2015).

27. M. L. Brewer, D. Dymock, R. L. Brady, B. B. Singer, M. Virji, D. J. Hill, Fusobacterium spp. target human CEACAM1 via the trimeric autotransporter adhesin CbpF. J. Oral Microbiol. 11, 1565043 (2019).

28. J. Bassler, B. Hernandez Alvarez, M. D. Hartmann, A. N. Lupas, A domain dictionary of trimeric autotransporter adhesins. Int. J. Med. Microbiol. 305, 265–275 (2015).

29. J. Strauss, G. G. Kaplan, P. L. Beck, K. Rioux, R. Panaccione, R. Devinney, T. Lynch, E. Allen-Vercoe, Invasive potential of gut mucosa-derived Fusobacterium nucleatum positively correlates with IBD status of the host. Inflamm. Bowel Dis. 17, 1971–1978 (2011).

30. Y. Yang, W. Weng, J. Peng, L. Hong, L. Yang, Y. Toiyama, R. Gao, M. Liu, M. Yin, C. Pan, H. Li, B. Guo, Q. Zhu, Q. Wei, M.-P. Moyer, P. Wang, S. Cai, A. Goel, H. Qin, Y. Ma, Fusobacterium nucleatum Increases Proliferation of Colorectal Cancer Cells and Tumor Development in Mice by Activating Toll-Like Receptor 4 Signaling to Nuclear Factor-κ, Up-Regulating Expression of MicroRNA-21. Gastroenterology (2016), doi:10.1053/j.gastro.2016.11.018.

31. M. Xu, M. Yamada, M. Li, H. Liu, S. G. Chen, Y. W. Han, FadA from Fusobacterium nucleatum utilizes both secreted and nonsecreted forms for functional oligomerization for attachment and invasion of host cells. J. Biol. Chem. 282, 25000–25009 (2007).

32. U. K. Gursoy, E. Könönen, V.-J. Uitto, Intracellular replication of fusobacteria requires new actin filament formation of epithelial cells. APMIS, 116, 1063–1070 (2008).

33. M. K. H. Schindler, M. S. Schütz, M. C. Mühlenkamp, S. H. M. Rooijakkers, T. Hallström, P. F. Zipfel, I. B. Autenrieth, Yersinia enterocolitica YadA mediates complement evasion by recruitment and inactivation of C3 products. J. Immunol. 189, 4900–4908 (2012).

34. J. Eitel, P. Dersch, The YadA Protein of Yersinia pseudotuberculosis Mediates High-Efficiency Uptake into Human Cells under Environmental Conditions in Which Invasin Is Repressed. Infect. Immun. 70, 4880–4891 (2002).

35. C. Wu, A. A. M. Al Mamun, T. T. Luong, B. Hu, J. Gu, J. H. Lee, M. D’Amore, A. Das, H. Ton-That, Forward Genetic Dissection of Biofilm Development by Fusobacterium nucleatum: Novel Functions of Cell Division Proteins FtsX and EnvC. MBio, 9 (2018), doi:10.1128/mBio.00360-18.

36. H. Nariya, S. Miyata, M. Suzuki, E. Tamai, A. Okabe, Development and application of a method for counterselectable in-frame deletion in Clostridium perfringens. Appl. Environ. Microbiol. 77, 1375–1382 (2011).

37. A. Ikegami, P. Chung, Y. W. Han, Complementation of the fadA mutation in Fusobacterium nucleatum demonstrates that the surface-exposed adhesin promotes cellular invasion and placental colonization. Infect. Immun. 77, 3075–3079 (2009).

38. A. Manson McGuire, K. Cochrane, A. D. Griggs, B. J. Haas, T. Abeel, Q. Zeng, J. B. Nice, H. MacDonald, B. W. Birren, B. W. Berger, E. Allen-Vercoe, A. M. Earl, Evolution of invasion in a diverse set of Fusobacterium species. MBio. 5, e01864 (2014).

39. I. D. Bobanga, F. Allen, N. R. Teich, A. Y. Huang, Chemokines CCL3 and CCL4 Differentially Recruit Lymphocytes in a Murine Model of Early Metastatic Colon Cancer. J. Surg. Res. 186, 515–516 (2014).

40. Y. Itatani, K. Kawada, S. Inamoto, T. Yamamoto, R. Ogawa, M. M. Taketo, Y. Sakai, The Role of Chemokines in Promoting Colorectal Cancer Invasion/Metastasis. Int. J. Mol. Sci. 17 (2016), doi:10.3390/ijms17050643.

41. V. Papayannopoulos, Neutrophil extracellular traps in immunity and disease. Nat. Rev. Immunol. 18, 134–147 (2018).

42. M. Doke, H. Fukamachi, H. Morisaki, T. Arimoto, H. Kataoka, H. Kuwata, Nucleases from Prevotella intermedia can degrade neutrophil extracellular traps. Mol. Oral Microbiol. 32, 288–300 (2017).

43. B. P. Lima, W. Shi, R. Lux, Identification and characterization of a novel Fusobacterium nucleatum adhesin involved in physical interaction and biofilm formation with Streptococcus gordonii. Microbiologyopen. 6 (2017), doi:10.1002/mbo3.444.

44. C. W. Kaplan, R. Lux, S. K. Haake, W. Shi, The Fusobacterium nucleatum outer membrane protein RadD is an arginine-inhibitable adhesin required for inter-species adherence and the structured architecture of multispecies biofilm. Mol. Microbiol. 71, 35–47 (2009).

45. C. W. Kaplan, X. Ma, A. Paranjpe, A. Jewett, R. Lux, S. Kinder-Haake, W. Shi, Fusobacterium nucleatum Outer Membrane Proteins Fap2 and RadD Induce Cell Death in Human Lymphocytes, doi:10.1128/IAI.00567-10.

46. Y. Ning, P. C. Manegold, Y. K. Hong, W. Zhang, A. Pohl, G. Lurje, T. Winder, D. Yang, M. J. LaBonte, P. M. Wilson, R. D. Ladner, H.-J. Lenz, Interleukin-8 is associated with proliferation, migration, angiogenesis and chemosensitivity in vitro and in vivo in colon cancer cell line models. Int. J. Cancer. 128, 2038–2049 (2011).

47. E. Cohen-Hillel, I. Yron, T. Meshel, A. Ben-Baruch, Interleukin 8 and cell migration to inflammatory sites: the regulation of focal adhesion kinase under conditions of migratory desensitization. Isr. Med. Assoc. J. 9, 579–583 (2007).

48. P.-L. Kuo, K.-H. ShenS H. Hung, Y.-L. Hsu, CXCL1/GROα increases cell migration and invasion of prostate cancer by decreasing fibulin-1 expression through NF-κB/HDAC1 epigenetic regulation. Carcinogenesis. 33, 2477–2487 (2012).

49. K. Mima, R. Nishihara, Z. R. Qian, Y. Cao, Y. Sukawa, J. A. Nowak, J. Yang, R. Dou, Y. Masugi, M. Song, A. D. Kostic, M. Giannakis, S. Bullman, D. A. Milner, H. Baba, E. L. Giovannucci, L. A. Garraway, G. J. Freeman, G. Dranoff, W. S. Garrett, C. Huttenhower, M. Meyerson, J. A. Meyerhardt, A. T. Chan, C. S. Fuchs, S. Ogino, Fusobacterium nucleatum in colorectal carcinoma tissue and patient prognosis. Gut (2015), doi:10.1136/gutjnl-2015-310101.

50. K. Nosho, Y. Sukawa, Y. Adachi, M. Ito, K. Mitsuhashi, H. Kurihara, S. Kanno, I. Yamamoto, K. Ishigami, H. Igarashi, R. Maruyama, K. Imai, H. Yamamoto, Y. Shinomura, Association of Fusobacterium nucleatum with immunity and molecular alterations in colorectal cancer. World J. Gastroenterol. 22, 557–566 (2016).

51. T. Yu, F. Guo, Y. Yu, T. Sun, D. Ma, J. Han, Y. Qian, I. Kryczek, D. Sun, N. Nagarsheth, Y. Chen, H. Chen, J. Hong, W. Zou, J.-Y. Fang, Fusobacterium nucleatum Promotes Chemoresistance to Colorectal Cancer by Modulating Autophagy. Cell. 170, 548–563.e16 (2017).

52. C. A. Brennan, W. S. Garrett, Gut Microbiota, Inflammation, and Colorectal Cancer. Annu. Rev. Microbiol. 70, 395–411 (2016).

53. A. D. Kostic, E. Chun, L. Robertson, J. N. Glickman, C. A. Gallini, M. Michaud, T. E. Clancy, D. C. Chung, P. Lochhead, G. L. Hold, E. M. El-Omar, D. Brenner, C. S. Fuchs, M. Meyerson, W. S. Garrett, Fusobacterium nucleatum potentiates intestinal tumorigenesis and modulates the tumor-immune microenvironment. Cell Host Microbe. 14, 207–215 (2013).

54. W. S. Garrett, The gut microbiota and colon cancer. Science? 364, 1133–1135 (2019).

55. R. Mizuno, K. Kawada, Y. Itatani, R. Ogawa, Y. Kiyasu, Y. Sakai, The Role of Tumor-Associated Neutrophils in Colorectal Cancer. Int. J. Mol. Sci. 20 (2019), doi:10.3390/ijms20030529.

56. J. Cools-Lartigue, J. Spicer, B. McDonald, S. Gowing, S. Chow, B. Giannias, F. Bourdeau, P. Kubes, L. Ferri, Neutrophil extracellular traps sequester circulating tumor cells and promote metastasis. J. Clin. Invest. (2013), doi:10.1172/JCI67484.

57. J. Park, R. W. Wysocki, Z. Amoozgar, L. Maiorino, M. R. Fein, J. Jorns, A. F. Schott, Y. Kinugasa-Katayama, Y. Lee, N. H. Won, E. S. Nakasone, S. A. Hearn, V. Küttner, J. Qiu, A. S. Almeida, N. Perurena, K. Kessenbrock, M. S. Goldberg, M. Egeblad, Cancer cells induce metastasis-supporting neutrophil extracellular DNA traps. Sci. Transl. Med. 8, 361ra138 (2016).

58. K. Mima, Y. Sukawa, R. Nishihara, Z. R. Qian, M. Yamauchi, K. Inamura, S. A. Kim, A. Masuda, J. A. Nowak, K. Nosho, A. D. Kostic, M. Giannakis, H. Watanabe, S. Bullman, D. A. Milner, C. C. Harris, E. Giovannucci, L. A. Garraway, G. J. Freeman, G. Dranoff, A. T. Chan, W. S. Garrett, C. Huttenhower, C. S. Fuchs, S. Ogino, Fusobacterium nucleatum and T Cells in Colorectal Carcinoma. JAMA Oncol. 1, 653–661 (2015).

59. R. Kalluri, The biology and function of fibroblasts in cancer. Nat. Rev. Cancer. 16, 582–598 (2016).

60. Y. Lin, J. Xu, H. Lan, Tumor-associated macrophages in tumor metastasis: biological roles and clinical therapeutic applications. J. Hematol. Oncol. 12, 76 (2019).

61. A. S. Abdulamir, R. R. Hafidh, L. K. Mahdi, T. Al-jeboori, F. Abubaker, Investigation into the controversial association of Streptococcus gallolyticus with colorectal cancer and adenoma. BMC Cancer. 9, 403 (2009).

62. J. H. Abed, J. Emgård, S. Chaushu, W. Garrett, G. Bachrach, Abstract 3300: Gal-GalNAc overexpressed in colorectal carcinoma mediates attachment and colonization of Fusobacterium nucleatum utilizing the Fap2 lectin. Cancer Res. 76, 3300–3300 (2016).

63. C. N. Spaulding, R. D. Klein, S. Ruer, A. L. Kau, H. L. Schreiber, Z. T. Cusumano, K. W. Dodson, J. S. Pinkner, D. H. Fremont, J. W. Janetka, H. Remaut, J. I. Gordon, S. J. Hultgren, Selective depletion of uropathogenic E. coli from the gut by a FimH antagonist. Nature. 546, 528–532 (2017).

64. L. Harbaum, M. J. Pollheimer, P. Kornprat, R. A. Lindtner, A. Schlemmer, P. Rehak, C. Langner, Keratin 7 expression in colorectal cancer--freak of nature or significant finding? Histopathology. 59, 225–234 (2011).

65. B. Huang, J. H. Song, Y. Cheng, J. M. Abraham, S. Ibrahim, Z. Sun, X. Ke, S. J. Meltzer, Long non-coding antisense RNA KRT7-AS is activated in gastric cancers and supports cancer cell progression by increasing KRT7 expression. Oncogene. 35, 4927–4936 (2016).

66. S. M. Singel, K. Batten, C. Cornelius, G. Jia, G. Fasciani, S. L. Barron, W. E. Wright, J. W. Shay, Receptor-interacting protein kinase 2 promotes triple-negative breast cancer cell migration and invasion via activation of nuclear factor-kappaB and c-Jun N-terminal kinase pathways. Breast Cancer Res. 16, R28 (2014).

67. J. Zheng, J. Meng, S. Zhao, R. Singh, W. Song, Campylobacter-induced interleukin-8 secretion in polarized human intestinal epithelial cells requires Campylobacter-secreted cytolethal distending toxin- and Toll-like receptor-mediated activation of NF-kappaB. Infect. Immun. 76, 4498–4508 (2008).

68. A. Kuznik, M. Bencina, U. Svajger, M. Jeras, B. Rozman, R. Jerala, Mechanism of endosomal TLR inhibition by antimalarial drugs and imidazoquinolines. J. Immunol. 186, 4794–4804 (2011).

69. W. S. 1. Rasband, ImageJ. Bethesda, MD: US National Institutes of Health. http://rsb.info.nih.gov/ij. 1997–2007 (1997).

